# Continuum dynamics and statistical correction of compositional heterogeneity in multivalent IDP oligomers resolved by single-particle EM

**DOI:** 10.1101/2020.06.16.154096

**Authors:** Barmak Mostofian, Russell McFarland, Aidan Estelle, Jesse Howe, Elisar Barbar, Steve L. Reichow, Daniel M. Zuckerman

## Abstract

Multivalent intrinsically disordered protein (IDP) complexes are prevalent in biology and control diverse cellular functions, including tuning levels of transcription, coordinating cell-signaling events, and regulating the assembly and disassembly of complex macromolecular architectures. These systems pose a significant challenge to structural investigation, due to the continuum dynamics imparted by the IDP and compositional heterogeneity resulting from characteristic low-affinity interactions. Traditional single-particle electron microscopy (EM) is a powerful tool for visualizing IDP complexes. However, the IDPs themselves are typically “invisible” by EM, undermining methods of image analysis and structural interpretation. To overcome these challenges, we developed a pipeline for automated analysis of common ‘beads-on-a-string’ type of assemblies, composed of IDPs bound at multivalent sites to the ubiquitous ~20 kDa cross-linking hub protein LC8. This approach quantifies conformational and compositional heterogeneity on a single-particle basis, and statistically corrects spurious observations arising from random proximity of bound and unbound LC8. After careful validation of the methodology, the approach was applied to the nuclear pore IDP Nup159 and the transcription factor ASCIZ. The analysis unveiled significant compositional and conformational diversity in both systems that could not be obtained from traditional single particle EM class-averaging strategies, and shed new light on how these architectural properties contribute to their physiological roles in supramolecular assembly and transcriptional regulation. Ultimately, we expect that this approach may be adopted to many other intrinsically disordered systems that have evaded traditional methods of structural characterization.

**Significance Statement:** Intrinsically disordered proteins (IDPs) or protein regions (IDRs) represent >30% of the human proteome, but mechanistically remain some of the most poorly understood classes of proteins in biology. This dearth in understanding stems from these very same intrinsic and dynamic properties, which make them difficult targets for quantitative and structural characterization. Here, we present an automated approach for extracting quantitative descriptions of conformational and compositional heterogeneity present in a common ‘beads-on-a-string’ type of multivalent IDP system from single-particle images in electron micrographs. This promising approach may be adopted to many other intrinsically disordered systems that have evaded traditional ensemble methods of characterization.

## INTRODUCTION

The role of intrinsically disordered proteins (IDPs) in organizing multivalent recruitment of regulatory proteins has been established in a wide range of systems, from metabolic enzymes, signal transduction scaffolds, kinases and gene regulation (1, 2). This range of functionality is made possible by the unique degree of conformational plasticity exhibited by IDP platforms that can be exploited for the recruitment of multiple binding partners with temporally regulated assembly. With the additional potential for tight control by post-translational modifications, IDP systems provide ideal substrates for their roles in cellular regulation (1, 3, 4). Despite the prevalence of IDPs (constituting as much as 1/3 of the human proteome; (5)) and their critical roles in cellular regulation, these systems remain some of the most mechanistically enigmatic and poorly understood components of molecular biology. This dearth in understanding stems from the very same intrinsic and dynamic properties that make multivalent IDP systems difficult targets for quantitative and structural characterization. The continuous and highly diverse conformational heterogeneity, in combination with often transient and/or multivalent binding properties that enable rapid and responsive regulatory roles, are notoriously difficult to characterize, as these features are often lost by traditional ensemble methods of structural characterization.

Our focus is on multivalent IDP strands which can form a duplex ladder-like assembly, reversibly cross-linked by the LC8 hub protein (DYNLL1) that form the ‘rungs’ of the ladder-like assembly (Fig. 1). Although the highly stable homodimer LC8 was originally characterized in complex with dynein (6, 7), a much broader role has now been well established with over 100 IDP binding partners in the cell (8, 9), impacting nucleopore assembly (10, 11), regulation of mitochondrial apoptosis (12), signal transduction (13), gene regulation (14–16) and many other processes (8, 17, 18). The complex binding and heterogeneity of LC8/IDP systems appears to be at the heart of its diverse functional roles: compositional heterogeneity is responsive to post-translational modifications and local LC8 concentration - *e.g.*, affecting spindle positioning in mitosis (19) and modular sensing for transcriptional activity (20), while large-scale conformational heterogeneity provides plasticity that is required for dynamic molecular machines, such as the dynein motor complex (21).

**Fig. 1:**
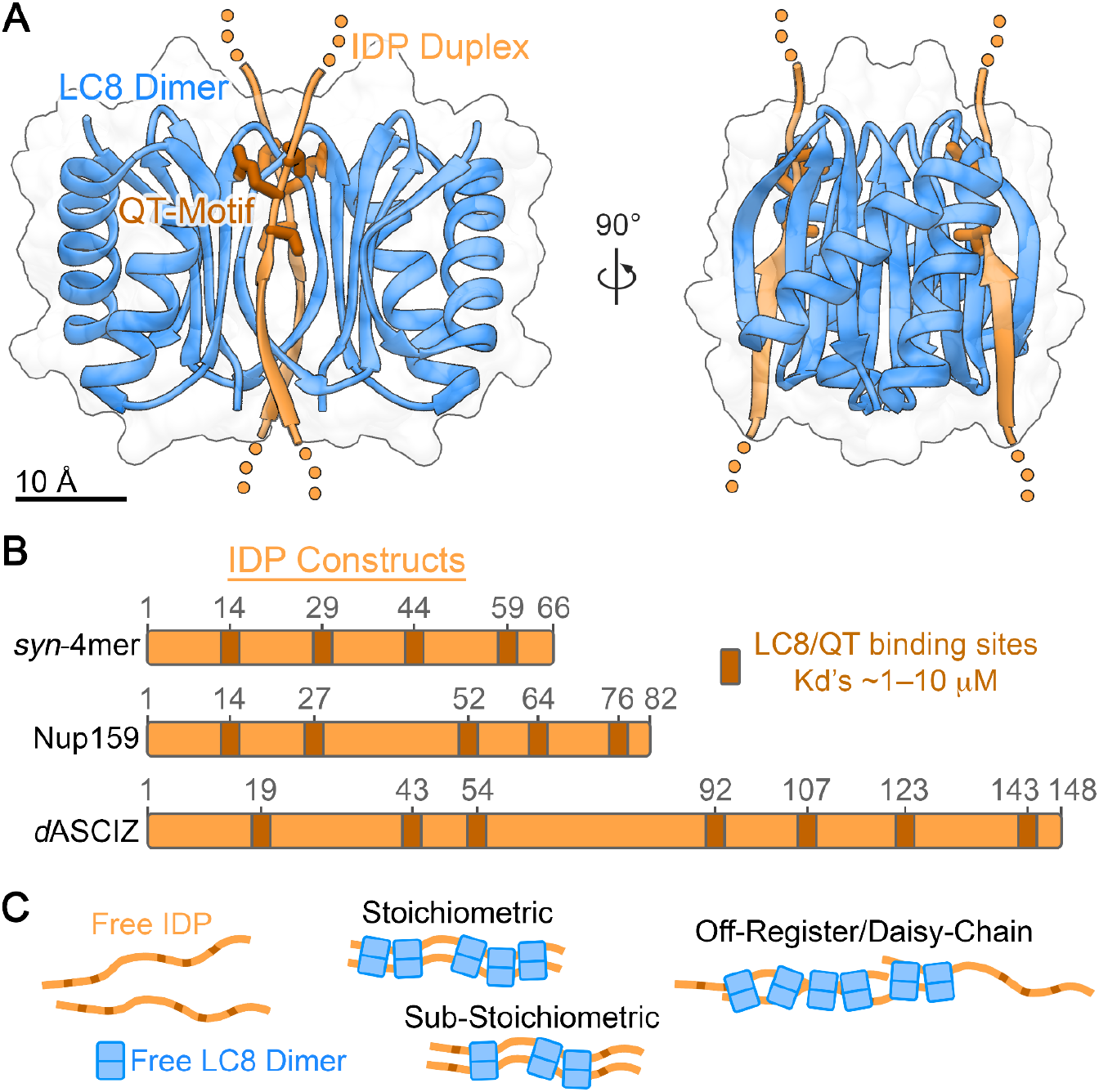
Overview of multivalent LC8/IDP systems. A) Crystallographic structure of the LC8 dimer (blue ribbon) bound to a duplex IDP (orange ribbon) (PDB ID 3GLW; (57)). Each LC8 protomer binds to a single IDP through a characteristic QT-motif (dark orange stick representation). Unresolved regions of the crystallized IDP construct are indicated by dotted line. B) Schematic representation of the IDP constructs under investigation, corresponding to the synthetic four-site IDP (*syn-*4mer), Nup159 and drosophila ASCIZ (*d*ASCIZ). LC8 binding QT-motifs are indicated in dark orange. Sequence numbering for each IDP construct and QT-motif is indicated. C) Illustration of various modes of assembly, ranging from (*left*) free IDP (orange) and free LC8 dimers (blue), to (*middle*) sub-stoichiometric and stoichiometric assemblies of LC8 bound to IDPs in a duplex fashion. In addition (*right)*, putative modes of off-register (or daisy-chain) assembly are illustrated, where > 2 IDPs are linked together by LC8 dimers.

Despite these established features of LC8/IDP systems, crucial structural and mechanistic knowledge gaps remain due to the inherent dynamical properties and transient formation of multiple oligomeric states that are key to their cellular function (22, 23). Notably, quantification of structural and compositional heterogeneity is lacking. Indeed, the dynamic nature of the disordered LC8/IDP complexes render structural determination by crystallography intractable. Aggregation, limited solubility, and conformational heterogeneity add to the challenges for characterization by NMR (24, 25). Previous analysis of the LC8/Nup159 system by single particle EM have been successful and visualizing the fully assembled oligomer, where five LC8 dimers appear uniformly arranged into a ladder-like assembly in two-dimensional class averages (26). However, the underlying complexity of conformational and configurational states present in this system were not characterized in this study.

Significant developments have been made in the field of high-resolution single-particle CryoEM image analysis for the characterization of conformational heterogeneity (*e.g.*, (27–33)). While extremely powerful, these approaches are most effective at characterizing large complexes (typically >100 kDa) needed to generate the necessary contrast in CryoEM images for accurate 3D alignment and are most effective at resolving discrete states of conformational heterogeneity. However, the small size of the LC8 dimer (~20 kDa), coupled with the broad continuum of conformational states in LC8/IDP complexes make these “beads-on-a-string” systems intractable to current high-resolution methods in CryoEM. Recently, we leveraged the high-contrast (and low-resolution) method of negative stain EM (NSEM) to directly visualize the highly heterogeneous multivalent LC8/IDP assemblies formed by the transcription factor ASCIZ, which regulates LC8’s own cellular concentration (22). However, this workflow was tremendously labor-intensive and subject to manual interpretation. Furthermore, quantifying the continuum of conformational states that abrogated the validity of traditional 2D classification results could not be readily assessed by such manual methods.

Here, we present an automated approach for single-particle *distribution* analysis, in contrast to standard single-particle class averaging that can suppress heterogeneity, for quantifying the conformational and compositional states of LC8/IDP ‘beads-on-a-string’ that are resolved in NSEM micrographs. To overcome potential artifacts arising due to low-affinity interaction that are characteristic of LC8/IDP systems (K_d_ ~1 – 10 μM), we apply a statistical correction process to estimate the effects caused by random proximity of free LC8 particles. The methodology was developed and validated using an artificial LC8/IDP system designed with 4 equivalent LC8 binding sites (termed *syn*-4mer), and then applied to two biological IDP systems, the nuclear pore protein Nup159 and the transcription factor ASCIZ (Fig. 1B). This approach recapitulated previous results for ASCIZ obtained by intensive manual methods, while further correcting over-estimation of small oligomeric species due to random proximity. For Nup159, we demonstrate a high-degree of conformational and configurational heterogeneity that had been obscured by previous ensemble methods of EM image analysis (26), and predict the presence of off-register type assemblies (Fig. 1C). Ultimately, we expect that this method may be generalized to obtain quantitative measurements on structure/assembly of many other ‘beads-on-a-string’ type multi-valent IDP systems.

## RESULTS

### Traditional characterization of a synthetic 4-site IDP construct by single-particle EM

To develop and validate our automated image analysis pipeline, we designed a synthetic 4-site LC8/IDP system (termed *syn*-4mer) (Fig. 1A,B). The QT-binding motif used in this IDP construct is based on a peptide sequence from the protein CHICA (34), selected for the reasonable binding affinity (K_d_ ~ 0.4 μM for the single site peptide). Each QT-motif is separated by a model flexible linker design (Methods). A tight LC8/*syn*-4mer interaction was validated by isothermal titration calorimetry (K_d_ of ~40 nM) and analytical ultra-centrifugation (AUC) (Supplemental Fig. 1). For initial structural characterization, the purified complex of LC8/*syn*-4mer was negatively stained and visualized by traditional single-particle EM image analysis (Fig. 2A). As expected under the dilute conditions required for NSEM (*i.e.*, below the K_d_ of 40 nM), a mixture of free LC8 dimers and assembled LC8/*syn*-4mer complexes were readily observed. Free LC8 dimers appear as small punctate densities (~5 nm diameter), while assembled oligomers appear as chains of 2 – 4 LC8’s separated by a characteristic spacing (4.7 ± 0.43 nm) dictated by the designed *syn*-4mer IDP.

**Fig. 2:**
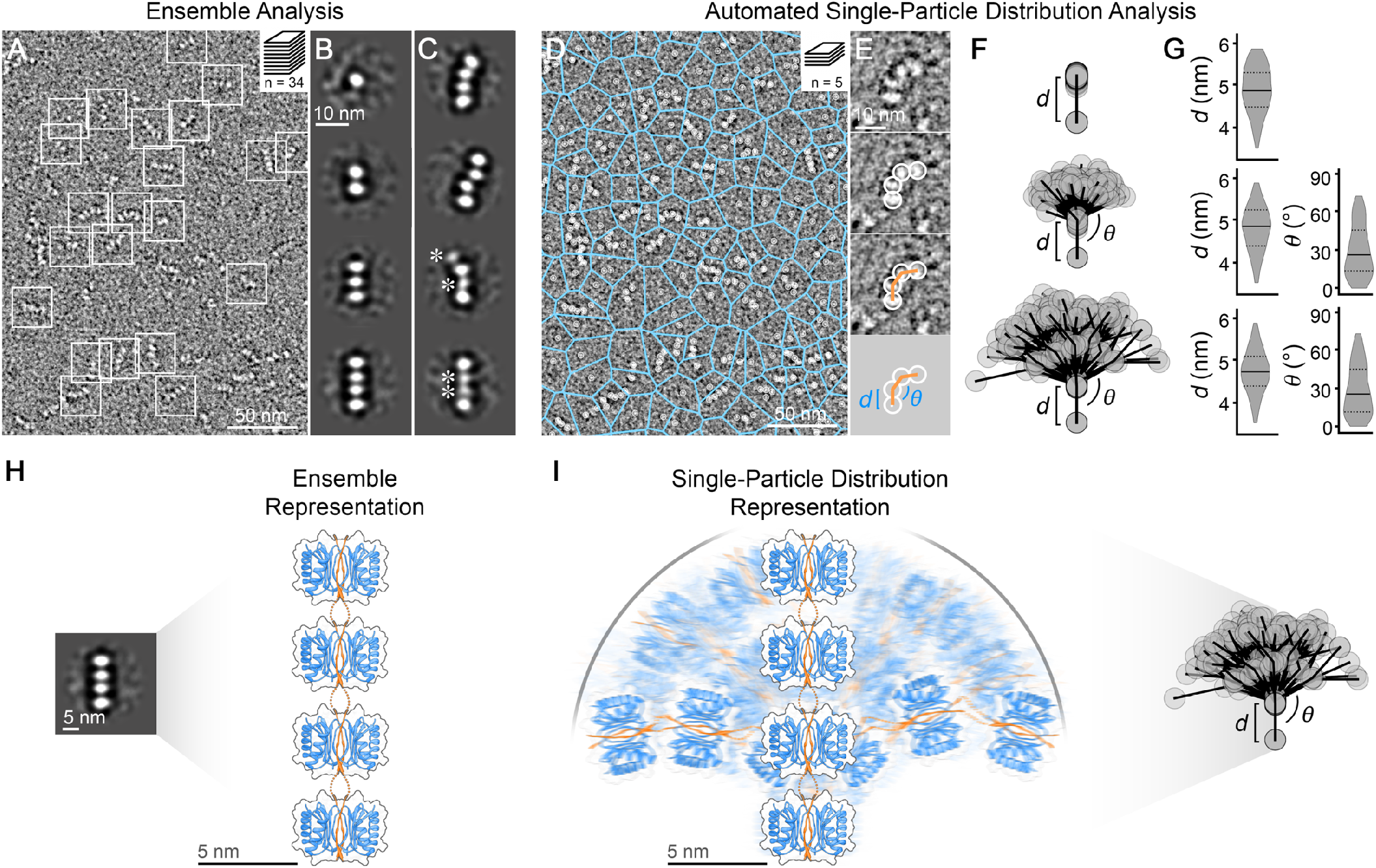
Ensemble versus automated single-particle characterization of LC8/IDP oligomers. A) Micrograph of negatively stained LC8/*syn*-4mer complexes. Representative complexes indicated by white box. Scale bar = 50 nm. *Inset*, indicates the number of micrographs collected for traditional ensemble analysis (n = 34) B) Representative two-dimensional (2D) class-averages depict free LC8 (*top*) and a range of assembled LC8/*syn*-4mer complexes (2-mers to 4-mers) present in the image dataset that were well-resolved. C) 2D class-averages of assembled LC8/*syn*-4mer complexes displaying varying degree of conformational heterogeneity. Asterisk indicate densities of LC8 that display blurred features that are less well-resolved, indicative of unresolved conformational/configurational heterogeneity. Scale bar = 10 nm in panels B and C. D) Micrograph of negatively stained LC8/*syn*-4mer complexes as shown in panel A, with auto-picked LC8 densities highlighted as white circles and the single-linkage clusters indicated with blue lines, calculated as edges from a Voronoi partitioning of cluster centers. Scale bar = 50 nm. *Inset*, indicates the number of micrographs collected for automated single-particle distribution analysis (n = 5) E) Zoom view from panel D, showing the automated classification and geometric analysis workflow. Individual LC8 dimers are selected in an automated fashion (white circles) and classified by our scoring function (represented by orange line). Geometric descriptions of the assigned oligomers are then extracted for analysis (*e.g.* LC8 to LC8 separation distances (*d*) and bend angles (*Θ*)). Scale bar = 10 nm. F) Illustrative representation of classified LC8/*syn*-4mer oligomers obtained by automated single-particle distribution analysis (n = 100 for each class of oligomer). Individual *n*-mers are aligned along the connection of the first two LC8 dimers (represented as grey circles). The conformational heterogeneity of identified oligomers is illustrated by the variability in separation distances (*d*) and bend angles (*Θ*). G) Violin plots showing the distribution of separation distances (*d*) and bend angles (*Θ*) of all 2-mers, 3-mers, and 4-mers. The solid line indicates the median and the dotted lines indicate the corresponding first and third quartiles. A full table of statistics is reported in Supplemental Table 2. H) Illustrative interpretation of *syn*-4mer duplex (orange ribbon) with four assembled LC8 dimers (blue) obtained by ensemble 2D classification methods. I) Illustrative interpretation of *syn*-4mer duplex (orange ribbon) with four assembled LC8 dimers (blue) obtained by automated single-particle distribution analysis, demonstrating the wide spectrum of conformational states accessible by the IDP scaffold.

A total of ~4150 LC8/*syn*-4mer oligomers and free LC8 particles were extracted from 34 micrographs, and subjected to traditional reference-free 2D classification procedures (Fig. 2B,C) (35). The most well resolved classes depict free LC8 and various LC8/*syn*-4mer assemblies, consisting of two, three or four LC8 dimers arranged in a nearly linear fashion (Fig. 2B). In addition to these ‘ideal’ classes, other conformational states are resolved, depicting arched and/or corrugated assemblies that are most recognizable in complexes with four bound LC8 dimers (Fig. 2C). Such variability is consistent with the range of conformational states expected to be accessible by the flexible linkers connecting neighboring LC8 binding sites. However, the degree of conformational heterogeneity resolved in 2D class averages appears to represent only a fraction of conformational states presented in the raw EM micrographs. The limitations of this ensemble approach is further apparent in several of the resulting 2D class averages, where LC8 densities often appear weak or blurred due to the underlying heterogeneity present in the images contributing to the ensemble representations (asterisks in Fig. 2C). Similar artifacts were present in our previously reported 2D class averages of LC8/ASCIZ system (22), and other IDP assemblies (36–38). Such artifacts are characteristic of EM 2D class averages where connected proteins, or domains, exhibit uncoupled and/or continuum dynamic behavior.

This analysis demonstrates that while the traditional approach of 2D class averaging significantly improves the overall signal-to-noise present in the raw images, the resulting average representations depict only a fraction of the underlying structural heterogeneity that can be observed at the single-particle level for such highly disordered beads-on-a-string type assemblies. Indeed, a majority of conformers that are part of the continuum of states are not represented by these results. While additional insights into the underlying conformational heterogeneity may be obtained by characterizing a much larger image dataset, such brute-force approaches would still be challenged by the continuum dynamics that are characteristic of IDPs.

### Automated single-particle distribution analysis resolves the continuum of conformational states in the LC8/*syn*-4mer system

The limitations of class averaging methods described above inspired the development of an automated image analysis pipeline that provides oligomer species populations and conformational distributions as assessed at the single-particle level. Our single-particle distribution analysis builds on two pillars: first, a straightforward, interpretable scoring function based on geometric and signal intensity criteria that is trained on a small set of manually selected oligomers; and second, a novel self-consistency analysis capable of correcting naïvely assigned oligomer populations based on the possibility of random proximity of oligomers and free LC8 particles.

To facilitate this approach, we treated each LC8 dimer (*a.k.a.* bead) independently, that is by not assuming the assembly state prior to analysis (Fig. 2D, white circles). The obtained coordinates of LC8 dimers were then subjected to single-linkage clustering. A scoring function was then applied to all possible oligomers within a cluster, with priority given to the largest possible oligomer that scored above a defined threshold. The score threshold was set to a low (permissive) value based on calibrated bead-to-bead distances and angles obtained from a small training dataset of hand-selected oligomers (Supplemental Fig. 4). Finally, a distance filtering step is applied to avoid assignments within crowded regions of the micrograph, by setting a minimum distance of 9 nm between assigned oligomers and other neighboring LC8 particles (Supplemental Fig. 5).

This approach was applied to a test dataset of ~17k isolated LC8 particles retrieved from only 5 micrographs (Fig. 2D-G). The output of this automated analysis provides a quantitative geometric description of the conformational state of each oligomer, defined by the center-to-center distance separating neighboring LC8 dimers (*d*), and the bend angle (θ) defined by three neighboring LC8 ‘beads’ (Fig. 2E-G). The resulting coordinates were plotted to visualize a representative ensemble of conformational states present in each oligomer class (Fig. 2F). The distribution of separation distances (*d*) and bend angles (θ) for each class was very similar, and consistent with the symmetrical design of the *syn-* 4mer IDP (Fig. 2G). In each class, the average separation distance (*d*) was equal to ~4.8 nm (± 0.5 nm), while the average bend angle (θ) was ~29° (± 20°).

The LC8-to-LC8 separation distances are consistent with those obtained in 2D class averages and with the length of the synthetic IDP, designed with 15 residues separating each QT recognition motif (Fig. 1B). A fully extended polypeptide of 15 residues would be expected to extend ~5.3 nm (*i.e.*, ~3.5 Å per residue), while a completely random polypeptide chain would be expected to follow a random-walk distribution, resulting in an average separation distance of ~1.4 nm (3.5 Å * √N_residues_ (39)). Thus, the center-to-center distance distribution obtained for the LC8/*syn*-4mer complexes suggests that the IDP adopts a primarily extended state, with only partial random character. Such characteristics are consistent with atomic models of LC8/IDP complexes where 10 amino acids of the QT recognition motif adopt an extended conformation when bound to LC8, but with a 5 residue linker between recognition motifs remaining flexible (Fig. 1A and Fig. 2H,I). This short flexible linkage is sufficient to facilitate the high degree of bend angles that lead to the continuum of conformational states presented in Fig. 2F,I, and minimum end-to-end distances between terminal LC8s that reach ~10 nm for the fully assembled 4-mer (Supplemental Fig. 4).

In comparison to the results obtained by traditional 2D class averaging methods, the single particle distribution approach applied here harvested a much greater degree of conformational states (distribution of bend angles), providing a much closer reflection of the heterogeneity observed in the raw micrographs. The conformational flexibility of this IDP complex is dramatized by a series of movies concatenating snapshots of oligomer images extracted from the micrographs (Supplemental Movies 1 and 2). This range of conformational motion would be difficult if not impossible to extract from class-averaged data. At the same time, geometric descriptions of the formed assemblies may be readily analyzed for quantitative measures and/or comparison between systems to assess the effects of LC8 assembly onto a variety of biological IDP scaffolds.

### Statistical correction for unbound LC8 particles refines species population profiles: 4-site system

An additional strength of our single-particle workflow is the ability to extract quantitative counts of identified species populations, which presents both new challenges and opportunities. Of particular interest is the question of whether off-register (or daisy-chain) type assembly occurs in LC8/IDP systems (Fig. 1C), which may be inferred by the presence of species with five or more assembled LC8 dimers in the *syn-*4mer dataset. However, oligomer assignments based purely on visible criteria such as geometry and signal intensity (labeled ‘initial’ assignments in Fig. 3), must be considered naïve because they cannot account for the possibility of random association among oligomer and/or free LC8 species. For example, a bonafide LC8/*syn*-4mer with four LC8’s randomly deposited on the EM grid in close proximity to a free LC8 dimer may be naïvely interpreted as evidence for off-register assembly because the IDP itself is not directly resolved by NSEM. Therefore, the invisibility of IDPs stringing together LC8 particles requires additional analysis of particle positions, extending ideas of correlation and Ripley’s K function analysis (40).

**Fig. 3:**
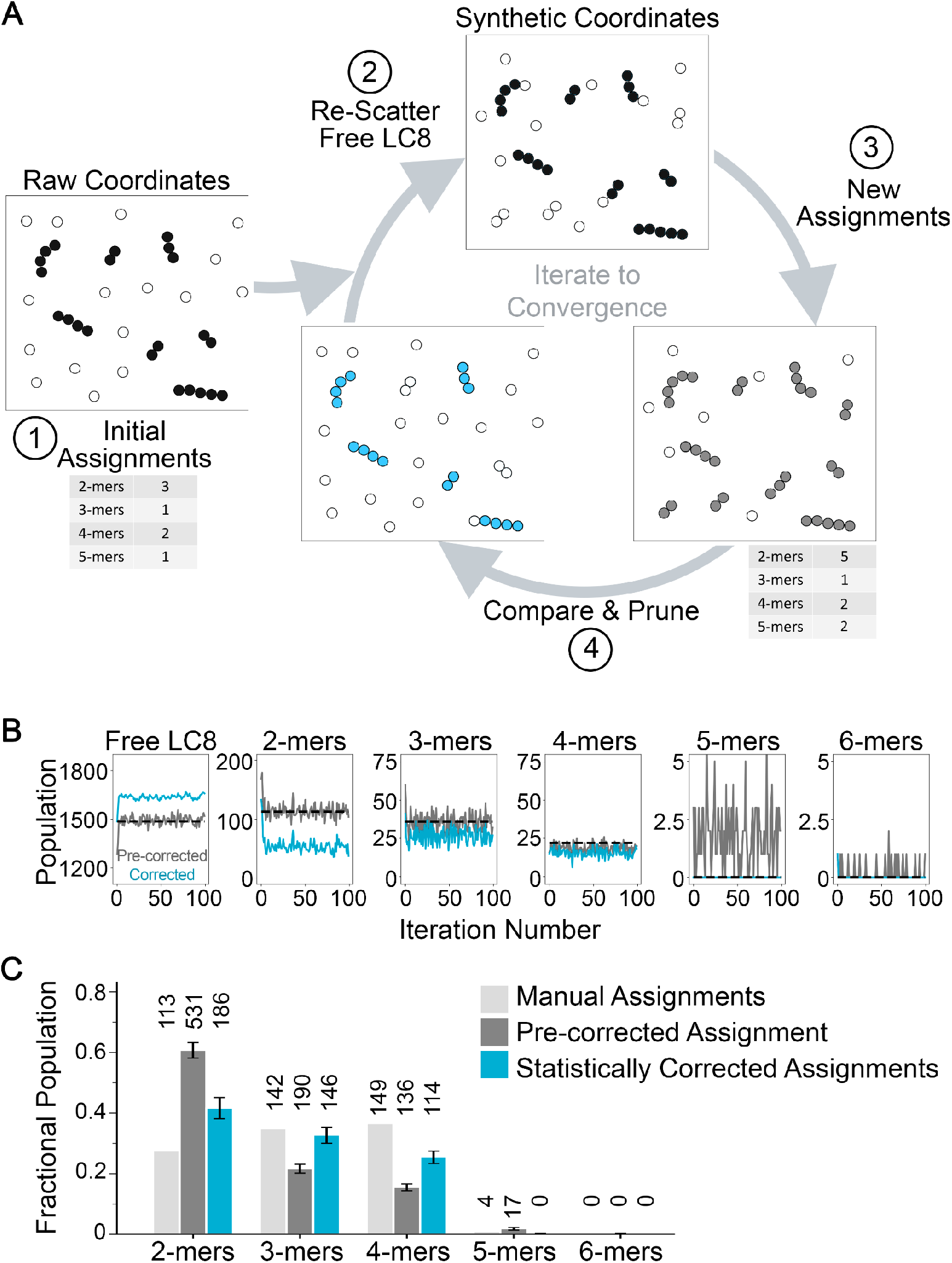
Self-consistent statistical correction of unbound LC8 particles. A) Initial oligomer assignments of the experimental micrograph are corrected to account for random proximity effects resulting from free LC8 particles. Accepting at first the initial oligomer assignments (black filled circles), the free LC8 population (white filled circles) is randomly rescattered to create a synthetic micrograph, from which new oligomer populations are obtained (grey filled circles). Deviations between pre-corrected and initial assignment populations are used to ‘correct’ the initial oligomer counts for the next round of rescattering (cyan filled circles). The rescattering and correction procedure is then iterated until self-consistency is obtained. B) The pre-corrected (gray) and corrected (cyan) populations of oligomers and free LC8 as a function of iteration number during the statistical correction simulation of one example micrograph with LC8 dimers bound to *syn*-4mer. At every iteration, the pre-corrected populations are from the synthetic micrograph and the corrected populations are the putative oligomers from the experimental micrograph after pruning. The dashed black lines indicate the corresponding populations of initial oligomer assignments in the experimental micrograph, to which the gray lines are expected to converge. Note that, in this example, the significantly overestimated number of 2-mers (and the vastly underestimated number of free LC8) is corrected within the first 10-20 iteration cycles. C) The fractional population distribution of LC8/*syn*-4mer oligomeric states obtained by manual inspection (light gray), by automated ‘pre-corrected’ single-particle distribution analysis (dark gray) and by statistical correction (blue). Total number of classified states are indicated above each bar. Error bars correspond to the effective standard deviation as described in Methods.

To provide an estimate for the actual number of the underlying oligomers, the experimental process of randomly depositing single LC8 particles onto the EM grid was iteratively simulated and the degree of artifactual oligomer creation was evaluated in order to obtain a self-consistent estimate of the true underlying oligomer populations (Fig. 3A). In every stage of the iterative process, synthetic micrographs are generated by randomly positioning the population of free LC8 particles while the positions of initially assigned oligomers remained fixed, and the resulting synthetic micrograph is re-classified and scored. This rescattering procedure inevitably alters the outcome of the classification process, as oligomers may be lengthened or created by random proximity of free LC8. By comparing these results to initial assignments, the putative list of oligomers can be iteratively refined, as detailed in Methods and Supplemental Fig. 6. Self-consistency is assessed by agreement of the pre-corrected populations from the synthetic micrographs (fluctuating gray lines in Fig. 3B) and the initial populations from the experimental micrograph (dashed black lines).

The results of the statistical correction analysis are presented in Fig. 3B,C. Populations of 3-mer and 4-mers are only mildly perturbed as compared to the naïve predictions; however, the population of 2-mers decreases dramatically, indicating a significant fraction of the original assignments reflected random proximity of free LC8 under these experimental conditions. Of greater interest, the analysis suggests that the population of 5-mers (*i.e.*, evidence of off-register/daisy-chain assembly) that were originally assigned are likely artifacts of random association between assembled 4-mers and free LC8 particles. In other words, we find it unlikely that off-register binding occurs in the 4-site system at the concentrations used for EM, fitting with the solution-state analysis by AUC that shows a homogeneous population assembly formed at much higher concentrations (Supplemental Fig. 1).

### Validation of the single-particle distribution analysis routine

The oligomers initially assigned by our scoring algorithm were validated by comparison to manually assigned oligomers in two phases: (*i*) assessment of whether the automated pipeline recovered oligomers assigned manually by the microscopist, and (*ii*) microscopist assessment of the quality of additional automated assignments not originally selected by the microscopist (Supplemental Fig. 3). Our assessment explicitly acknowledges that we lack “ground truth” oligomer assignments because it is impossible to unambiguously distinguish background from noise, or to distinguish visually between random proximity and true oligomerization. The latter point, in fact, motivated the statistical correction analysis (described in the previous section). However, because our correction procedure is based on statistical inferences and not on observable structural features, the results of this approach could not be validated in a similar way.

The first validation analysis revealed that the automated pipeline recovered ~90% or more of manually assigned 2-mer, 3-mer, and 4-mer oligomers (Supplemental Fig. 4). A lower proportion (~75%) of manually assigned 5-mers were recovered, but as shown (Fig. 3), it is likely that the naïvely assigned 5-mers are the result of random association of smaller oligomers with free LC8s.

In addition, our automated analysis revealed the presence of oligomers beyond those identified by the microscopist. The second phase of validation revealed a range of phenomena that could be attributed to the discrepancy between microscopist-identified complexes and the automated procedure (Fig. 4). Most notably, among the discovered complexes (*i.e.*, those not originally identified by the microscopist), 40% or more were found by the microscopist to be acceptable upon inspection (Fig. 4A,B), with the remainder judged to be invalid (Fig. 4C,D). The invalid complexes, in turn, were approximately evenly split into two cases: in the first group, some of the assigned LC8 density(s) were judged to be too ambiguous to permit confident identification of an oligomer (*i.e.*, containing poorly resolved or weak LC8 particle density) (Fig. 4C). The second group of invalid assignments was characterized by overall unconvincing picks of single LC8 particles (*i.e.*, deemed to be falsely picked particles that were not clearly distinguishable from background noise) (Fig. 4D).

**Fig. 4:**
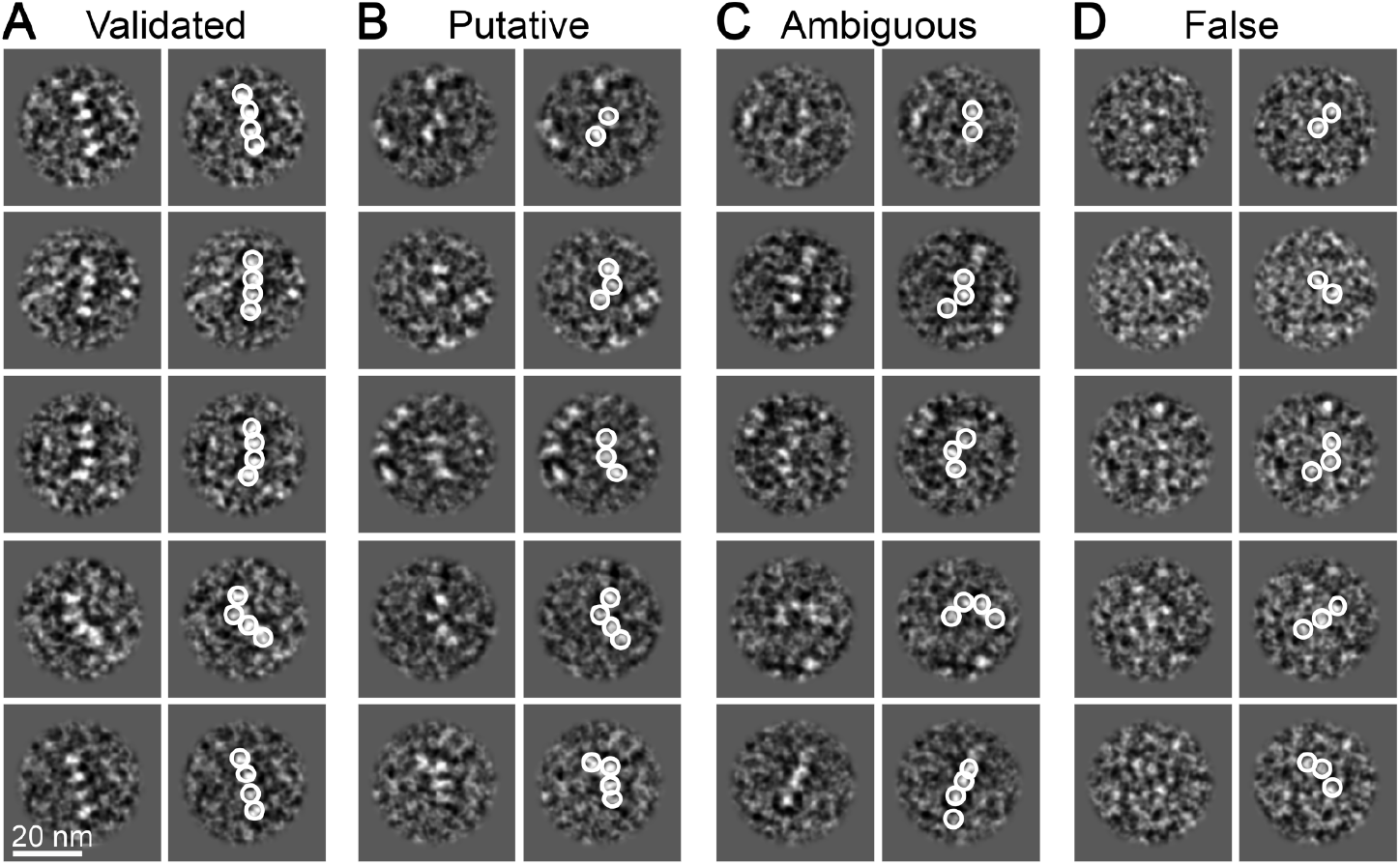
Validation of automated LC8/IDP oligomer assignments. A) Representative images of validated LC8/*syn*-4mer oligomers that were assigned by both the microscopists and the scoring function. B) Representative images of putative complexes assigned by scoring function and deemed to be acceptable by the microscopist upon re-evaluation. C) Representative images of complexes assigned by scoring function and deemed to be too ambiguous to confidently assign by the microscopists upon re-evaluation (*e.g.*, containing weak LC8 density and/or neighboring LC8 densities that were not autopicked). D) Representative images of complexes assigned by scoring function and deemed to be too falsely assigned by the microscopist upon re-evaluation (*e.g.*, containing autopicked densities corresponding to background carbon). Scale bar for all panels = 20 nm. In all panels, raw images are shown in the left column and autopicked results obtained by DoG Picker (55) shown by white circles.

Hence, it appears that most automated oligomer assignments that were deemed by the microscopists to be erroneous stemmed from unreliable particle picks, rather than from intrinsic problems with the oligomer identification algorithm. Although significant care was taken to optimize the automated particle picking parameters used in this study, spurious background picks were unavoidable. Indeed, the identification of such small ~20 kDa particles in NSEM micrographs is often ambiguous via manual inspection as well, and the automated approach we employed compared favorably to a variety of established particle picking tools (see Methods). Importantly, the population of spurious background picks leading to erroneous assignments is relatively small and may be further filtered by selecting a more stringent scoring threshold; the particle picker is a fully modular component of our pipeline.

### Single-particle distribution analysis of LC8/Nup159

With the protocol for single-particle distribution analysis validated using the LC8/*syn*-4mer system, we went on to examine more heterogeneous and biologically relevant LC8/IDP assemblies. The first case was the nuclear pore protein Nup159, an IDP which binds up to 5 LC8 dimers in a duplex fashion (Fig. 1) (11, 26). In contrast to the LC8/*syn*-4mer system, LC8/Nup159 displays a high-degree of compositional heterogeneity even under saturating conditions, as assessed by AUC (Supplemental Fig. 1). For our comparative analysis by NSEM, we performed both traditional 2D class-averaging and the single-particle distribution analysis protocol on a dataset obtained from 30 micrographs (Fig. 5 and Supplemental Fig. 7). Scoring and distance thresholds were defined as described in Methods and Supplemental Fig. 8.

**Fig. 5:**
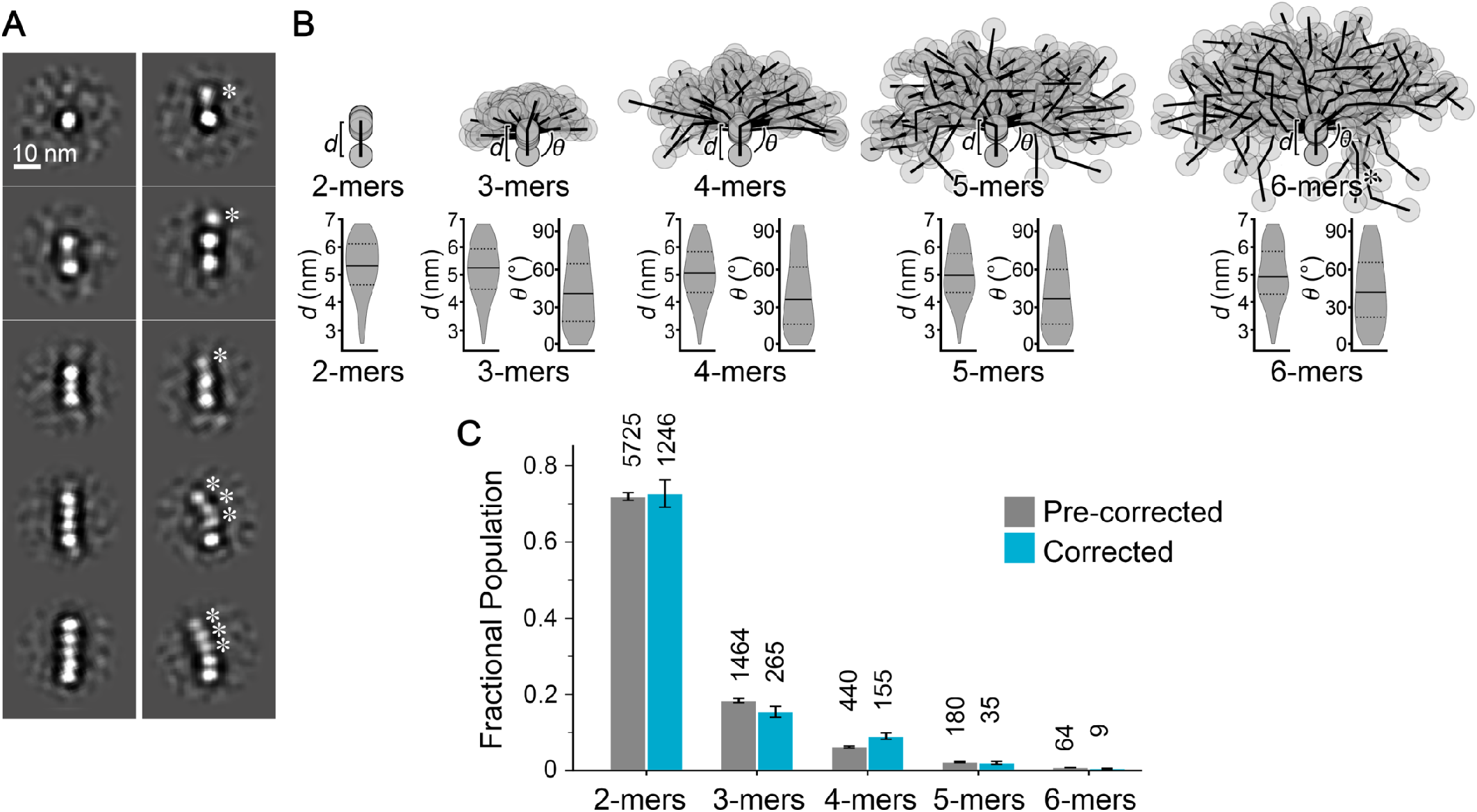
Quantitative characterization and statistical correction of LC8/Nup159 complex assemblies. A) Representative 2D class-averages depicting, (*left column*) free LC8 and a range of assembled LC8/Nup159 complexes ranging from 2-mers to 5-mers present in the image dataset that were well-resolved, and (*right column*) complexes displaying varying degrees of conformational heterogeneity. Asterisk indicate densities of LC8 that display blurred features that are less well-resolved, indicative of unresolved conformational/configurational heterogeneity. Scale bar = 10 nm. B) Illustrative representation of classified LC8/Nup159 oligomers obtained by automated single-particle distribution analysis (n = 100 for each class of oligomer). Individual *n*-mers are aligned along the connection of the first two LC8 dimers (represented as grey circles). The conformational heterogeneity is illustrated by the violin plots showing the distribution of separation distances (*d*) and bend angles (*Θ*) of all oligomers identified from a total of 30 micrographs. The solid line indicates the corresponding median and the dotted lines indicate the corresponding first and third quartiles. A full table of statistics is reported in Supplemental Table 2. C) The fractional population distribution of oligomeric states obtained by automated single-particle distribution analysis prior to correction (gray) and following statistical correction (blue). Total number of classified states are indicated above each bar. Error bars correspond to the effective standard deviation as described in Methods.

Traditional 2D classification analysis of ~5875 identified oligomers resolved a range of species, representing free LC8 dimers to fully assembled LC8/Nup159 complexes with 5 bound LC8 dimers (Fig. 5A and Supplemental Fig. 7). The fully-assembled complex appears similar to what was previously described by Stelter et al (26), with five LC8 dimers arranged in an ordered and nearly linear fashion, with each LC8 dimer separated by 4.5 ± 0.66 nm. However, in addition to these well-resolved oligomeric assemblies, many of the other 2D class averages demonstrate the presence of underlying conformational heterogeneity, resulting in the appearance of blurred LC8 densities and arranged in a non-linear fashion (Fig. 5A, asterisk and Supplemental Fig. 7).

The results of our automated single-particle distribution analysis once again revealed the extent of significant conformational heterogeneity that is apparent in the raw micrograph images (Fig. 5B and Supplemental Fig. 7). The quantified separation distances between LC8 particles were similar among the classified oligomer states, ranging from 5.0 ± 1.0 to 5.28 ± 1.0. Again, such distances are consistent with the length of linkers separating the 10-residue long QT-motifs (2 – 15 residue linkers, see Fig. 1) and the expected percentage of disordered versus structured IDP character that is induced by LC8 binding. The longer linker region within the Nup159 IDP (separating motifs 2 and 3), as compared to the *syn*-4mer, appear to result in a broader distribution of bend angles, which range from 39° ± 26 to 44° ± 27 in the Nup159/IDP assemblies (Fig. 5B). The result of this flexibility is an apparent continuous ensemble of conformational states supported by the IDP scaffold, and in the distribution of end-to-end distances measured between terminal LC8 dimers (Supplemental Fig. 8).

The population counts of oligomeric assemblies decay significantly with increasing size, with the 2-mer state being the most populated (Fig. 5C). Such behavior is expected for the moderate binding affinity between LC8 and the Nup159 IDP (K_d_ ~3 μM) and the sample dilution to nanomolar level (41). In addition to the expected stoichiometries of 2 – 5-mers, the initial automated assignment of oligomeric states also identified a significant population of 6-mers (*i.e.*, Nup159 bound to 6 LC8 dimers) (Fig. 5B), again indicating the possible existence of off-register/daisy-chain type assemblies even at these low concentrations (see Fig. 1). Remarkably, application of our statistical correction protocol does not rule out the existence of the off-register 6-mer species in this system (Fig. 5C and Supplemental Fig. 8). The significance of this intriguing finding is further discussed below. On the other hand, similar to the LC8/*syn*-4mer system, the corrected population of 2-mers is significantly reduced from the initial population based only on visible features. Indeed, the formation of randomly proximal 2-mers is expected whenever there is a substantial population of free LC8 particles. In contrast to the *syn*-4mer dataset, however, the relative population of 2-mers is not significantly altered following our statistical correction protocol (Fig. 5C).

### Single-particle distribution analysis of LC8/*d*ASCIZ

To further assess the effectiveness of our heterogeneity analysis, we characterized the transcription factor ASCIZ, which regulates synthesis of the LC8 protein to which it binds in a multivalent fashion. Drosophila ASCIZ (*d*ASCIZ) has seven LC8 binding sites (Fig. 1). We have shown that the LC8/*d*ASCIZ assembly exhibits significant compositional and conformational heterogeneity by NSEM, NMR and analytical ultra-centrifugation (22) and by native mass spectrometry (23). Manual analysis of the NSEM data was used to obtain a quantitative assessment of oligomer populations, and while conformational heterogeneity could be deduced from the raw micrograph images, a quantitative procedure to characterize these states was not practical.

To facilitate comparison of results to the *syn-*4mer and Nup159 systems described here, we re-analyzed the LC8/*d*ASCIZ NSEM dataset using the same workflow, as described in Methods (Fig. 6 and Supplemental Figs. 9 and 10). Traditional 2D classification analysis appeared to resolve only a subset of oligomeric states (Fig. 6A and Supplemental Fig. 9), while fully assembled 7-mers were not identified under the dilute conditions required for NSEM (Fig. 6C). These results are consistent with our previous analysis, and the characterized negative cooperativity that is present in this system (22). Furthermore, the same artifacts described for the *syn-*4mer and Nup159 systems resulting from the underlying conformational heterogeneity of this system are readily identified in the results of 2D class averaging (Fig. 6A, asterisks and Supplemental Figs. 8), and as previously described (22).

**Fig. 6:**
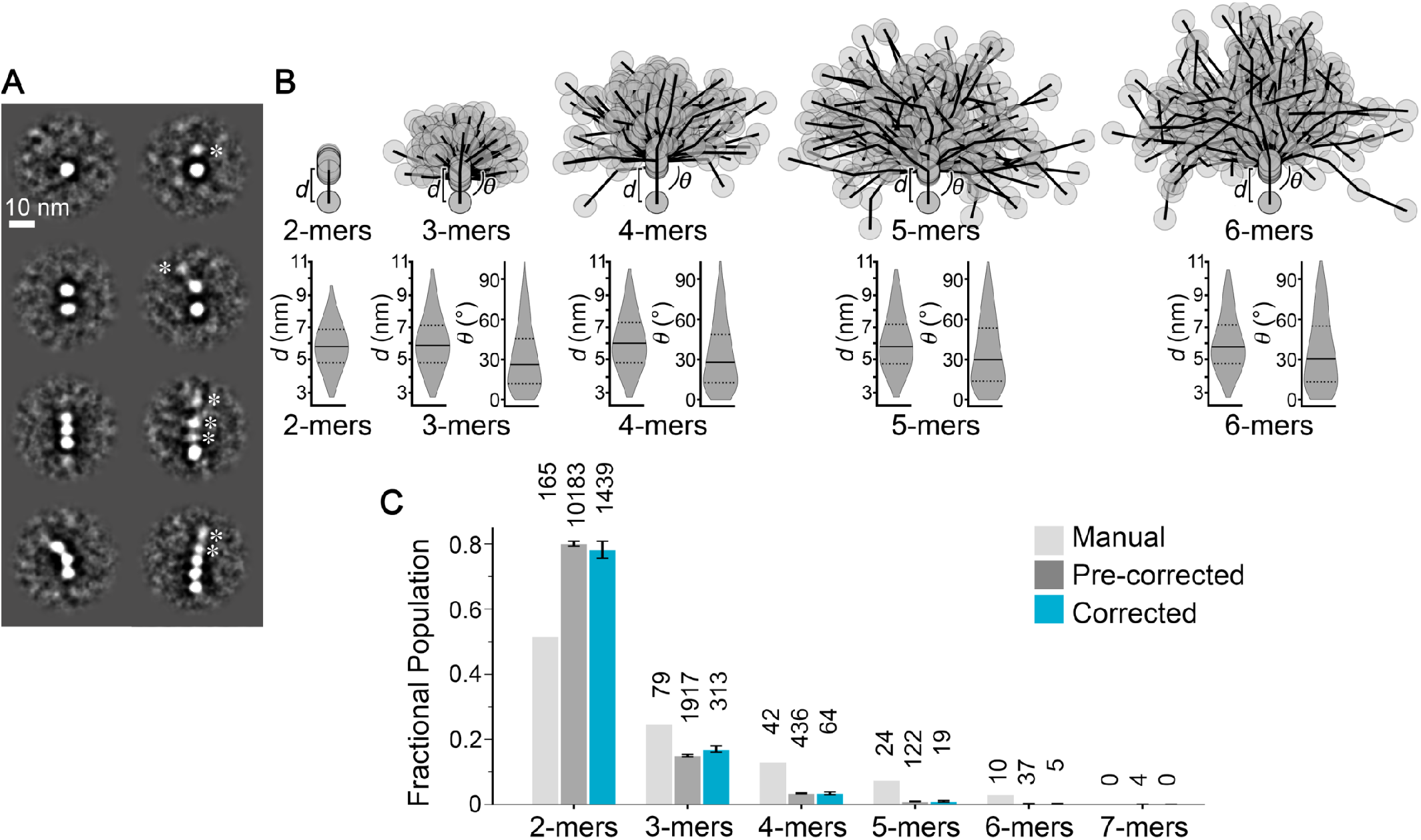
Quantitative characterization and statistical correction of LC8/*d*ASCIZ complex assemblies. A) Representative 2D class-averages depicting, (*left column*) free LC8 and a range of assembled LC8/*d*ASCIZ complexes ranging from 2-mers to 3-mers present in the image dataset that were well-resolved, and (*right column*) complexes displaying varying degree of heterogeneity. Asterisk indicate densities of LC8 that display blurred features that are less well-resolved, indicative of unresolved conformational/configurational heterogeneity. Scale bar = 10 nm. B) Illustrative representation of classified LC8/*d*ASCIZ oligomers obtained by automated single-particle distribution analysis (n = 100 for each class of oligomer, except for the 6-mer class which was limited to n = 37). Individual *n*-mers are aligned along the connection of the first two LC8 dimers (represented as grey circles). The conformational heterogeneity is illustrated by the violin plots showing the distribution of separation distances (*d*) and bend angles (*Θ*) of all oligomers identified from a total of 302 micrographs. The solid line indicates the corresponding median and the dotted lines indicate the corresponding first and third quartiles. A full table of statistics is reported in Supplemental Table 2. C) The fractional population distribution of oligomeric states obtained by manual inspection (light gray), automated single-particle distribution analysis prior to correction (dark gray) and following statistical correction (blue). Total number of classified states are indicated above each bar. Error bars correspond to the effective standard deviation as described in Methods. The populations obtained by manual inspection are based on a subset of only 50 micrographs.

Our single-particle distribution analysis once again provides a much more detailed and quantitative picture of the underlying conformational and compositional heterogeneity present in the LC8/*d*ASCIZ system. Due to complications associated with the extreme level of heterogeneity in this sample, our prior manual analysis was limited to < 350 total oligomers (extracted from ~300 micrographs) (22), whereas the present automated pipeline yielded more than an order of magnitude more oligomers from the same set of micrographs (Fig. 6C). Geometric distributions of LC8 bound to *d*ASCIZ portray a system with significant structural variability (Fig. 6B). Remarkably, despite the presence of a variety of linker lengths separating the 10-residue QT-motifs (ranging from 1 – 28 flexible residues), separation distances between bound LC8’s are similar to the *syn*-4mer and Nup159 assemblies, ranging from 5.8 ± 1.4 to 6.1 ± 1.7 nm (Fig. 6B). The end-to-end distance of terminal LC8 dimers becomes widely distributed with increasing valency, and reaches a minimum of ~10 nm for assembled 6-mers (Supplemental Fig. 10). This continuum of states is facilitated by the accessible bend angles, which are slightly more narrowly distributed as compared to Nup159, and more similar to the *syn-*4mer assemblies, ranging from 30° ± 23 to 36° ± 26 (Fig. 6B). The mechanistic explanation for this behavior is not yet clear, but might reflect some intrinsic behavior or communication between LC8 binding sites that is responsible for the characterized negative cooperativity displayed by *d*ASCIZ.

Population profiles of the assembled oligomeric states determined by manual assessment, by the initial automated process, and after statistical correction are in rough agreement (Fig. 6C). Notably, neither the manual nor the statistically corrected counts tally any fully saturated 7-mers, let alone larger potentially off-register species. In all cases, the 2-mer population is the most abundant species, representing almost 80% of the population by our automated workflow, while 3-mers and 4-mers make up the majority of other species detected. This assessment is consistent with results for 2D class averages, where well-resolved classes corresponded to only the 2-mer and 3mer populations (Fig. 6A), lending further validity to the results of the presented automated approach and statistical correction process.

## DISCUSSION

The continuum dynamics (conformational fluctuations) of IDP systems makes them generally difficult to structurally characterize with precision, and the emerging class of semi-ordered beads-on-a-string systems, such as the LC8-organized systems studied here, compound the challenges. The LC8 dimers ‘beads’ are too small for current high-resolution CryoEM methods, forcing the use of NSEM in this work where the IDPs themselves are not directly detectable. Adding to these challenges, multivalent LC8/IDP systems exhibit substantial compositional heterogeneity, originating from their relatively low binding affinities. Class-averaging methods common in EM analysis are inappropriate for such systems because they suppress heterogeneity by construction. We have therefore developed a single-particle distribution analysis pipeline. Such tools are expected to become increasingly valuable given that the multivalent systems studied here appear to exploit weak, reversible, multivalent binding in support of diverse nano-architectural and sensory roles, with the full scope of functions still being revealed (42–45).

The idea to extract single-particle information directly from individual electron micrograph images is not new (22, 36–38), but here we exploit geometric characteristics of polymeric systems to create an intuitive and reliable automated approach applicable to the growing, important class of beads-on-string systems (46–48). We show that the “functional form” of polymeric conformational properties in terms of bead-bead distances and three-bead angles, enabled adequate training based on only a few dozen manually picked oligomers. The automated approach then generates thousands of candidate structures that represented the full breath of conformational states observed in the single-particle population, which can be analyzed and/or filtered as needed. The statistical correction process allowed the microscopists to gain a more faithful visualization of the underlying compositional heterogeneity that may be obscured by large populations of unbound ligand/proteins. In this way, we were able to address possible off-register binding (Fig. 1), an emerging phenomenon (23). For example, for Nup159, it is possible that off-register binding is required to bridge the IDP dimers into the 8-fold symmetry of the nuclear pore complex and stabilize the higher order assembly (49) – providing an intriguing basis for future investigation. However, for other LC8/IDP systems, the potential for such types of off-register assembly may require suppression for LC8 to orchestrate its physiological roles.

The pipeline presented here can be improved in future studies. Better optimized single-particle picking algorithms would enhance oligomer assignment quality most directly, though we note that preliminary testing of some modern particle picking platforms (35, 50, 51) yielded sub-optimal results, presumably because of the small size and close proximity of LC8 particles. Technical aspects of the scoring and statistical correction procedures might also be improved, such as giving preference to pruning low-scoring oligomers/particles as part of the statistical correction procedure. The oligomer scoring function could also include the possibility of LC8 “beads” appearing in a non-sequential fashion, *e.g.*, accounting for cases where an LC8 dimer is missing in the middle of an oligomer. At the same time, it may be possible in some systems to correlate the extracted values of bead-to-bead distances with the known lengths of the IDP linkers, allowing identification of occupied binding sites.

Importantly, the methods developed here are readily extensible to other heterogeneous and dynamic multivalent IDP systems. It should be straightforward to generate synthetic micrographs for self-consistent analysis within the statistical correction procedure demonstrated here. Likewise, determining and scoring geometric and intensity criteria should also present few obstacles for other systems.

## METHODS

### Design of a synthetic 4-site LC8-binding protein

For algorithm development and training, we designed a novel LC8-binding peptide (termed *syn*-4mer) using a series of 4 repeats of the amino acid sequence RKAIDAATQTE, taken from the tight-binding LC8 motif of the protein CHICA (Uniprot Q9H4H8), which has a 0.4 μM affinity to LC8, making it one of the tightest-known LC8-binding motifs (34). The motif is spaced by uniform disordered linker sequences, totaling 3 linkers, and flanking GSYGS sequences were added to the N- and C-termini of the constructs to allow for quantification by absorbance at 280 nm. The final sequence is: GSYGSR**KAIDAATQTE**PKETR**KAIDAATQTE**PKETR**KAIDAATQTE**PKETR**KAIDAATQTE**GSYGS. In bold is the 10 amino acid segment that packs as a beta strand when bound to LC8.

### Protein expression and purification

A gene sequence for the LC8-binding *syn*-4mer peptide was purchased as a block (integrated DNA technologies, Coralville, Iowa) and cloned into a pET24d expression vector with an N-terminal Hisx6 affinity tag and a tobacco etch virus protease cleavable site. The protein was expressed in ZYM-5052 (52) auto-induction media at 37° C for 24 hr. Cells were harvested, lysed by sonication and purified in denaturing buffers containing 6 M urea on TALON resin. The 4-mer was dialyzed into non-denaturing buffer (25 mM tris pH 7.5, 150 mM NaCl) and further purified by gel filtration on a Superdex 75 Hi-load column (GE Health), in the same buffer. Domains of yeast Nup159 (residues 1096 – 1178) and drosophila ASCIZ (*d*ASCIZ, residues 241 – 388), full length LC8 of Saccharomyces cerevisiae and LC8 of Drosophila melanogaster were all expressed and purified as previously described (22, 34). All proteins were stored at 4° C and used within one week of purification.

### SEC-MALS

Size-exclusion chromatography (SEC) coupled to a multiangle light scattering (MALS) instrument was performed using an analytical SEC column of Superdex S200 resin (GE Healthcare) on an AKTA-FPLC (GE Healthcare), then routed through a DAWN multiple-angle light scattering and Optilab refractive index system (Wyatt Technology). The column was equilibrated to a buffer of 25 mM tris (pH 7.5), 150 mM NaCl, and 5 mM BME, then injected with 100 μL of LC8/*syn*-4mer complex in the same buffer at an estimated 2 μM particle concentration (16 μM LC8 + 4 μM *syn*-4mer, assuming 2:8 binding stoichiometry). We estimated the molar mass using the ASTRA software package, with a Zimm scattering model.

### Isothermal Titration Calorimetry

Isothermal titration calorimetry was carried out at 25°C using a VP-ITC microcalorimeter (Microcal) in a buffer of 25 mM tris (pH 7.5), 150 mM Nacl and 5 mM BME. A cell containing 9 μM syn-4mer was titrated with a solution of 300 μM LC8, across 32 injections of 8 μL. Peaks were integrated and fit to a single-site binding model in Origin 7.0.

### Analytical Ultracentrifugation

Samples of the *syn*-4mer peptide in complex with LC8 were prepared for sedimentation velocity analytical ultracentrifugation (SV-AUC) by mixing excess (8:1) LC8 with *syn*-4mer, then purifying the complex by gel filtration on a Superdex 200 column in a buffer of 25 mM tris (pH 7.5), 150 mM NaCl, and 5 mM β-mercaptoethanol. The estimated concentration of the syn-4mer/LC8 complex applied to SV-AUC was at a 4:1 ratio of syn-4mer (13.8 μM) and LC8 (55μM). The SV-AUC titration of LC8 into Nup159 was performed by mixing Nup159 (12.5 μM) and LC8 at LC8:Nup159 ratios of 0.5:1 to 8:1 in a buffer of 50 mM sodium phosphate (pH 7.5), 50 mM NaCl, 5 mM TCEP and 1 mM sodium azide. SV-AUC was performed on a Beckman Coulter Optima XL-A ultracentrifuge, equipped with optics for absorbance. Complexes were loaded into two-channel sectored centerpieces with a 12-mm path length and centrifuged at 42,000 rpm and 20° C. We collected 300 scans at 280 nm with no interscan delay, and fit data to a c(S) distribution using SEDFIT (53). Buffer density was calculated using Sednterp (54).

### LC8-IDP complex preparation for EM

LC8/syn-4mer complexes were prepared for electron microscopy studies by mixing excess (8:1) of the purified LC8 with the syn-4mer peptide and purifying the complexes by size-exclusion chromatography (SEC; Superdex 200, in a buffer of 25 mM tris pH 7.5, 150 mM NaCl and 5 mM BME). The Nup159 complex was formed by mixing equimolar amounts of LC8 and Nup159, without further purification. Negative stain EM grids were prepared by diluting the LC8 complexes to a final particle concentration of 16 nM (presumed to be fully bound complexes) in SEC buffer. A 3 μl drop of sample was applied to a glow-discharged continuous carbon coated EM specimen grid (400 mesh Cu grid, Ted Pella, Redding, CA). Excess protein was removed by blotting with filter paper and washing the grid two times with dilution buffer. The specimen was then stained with freshly prepared 0.75% (wt vol^-1^) uranyl formate (SPI-Chem).

### Electron microscopy

Negatively stained specimens were imaged on a 120 kV TEM (iCorr, FEI) at a nominal magnification of 49,000x at the specimen level. Digital micrographs were recorded on a 2K × 2K CCD camera (FEI Eagle) with a calibrated pixel size of 4.37 Å pixel^-1^ and targeted a defocus of 1.5 – 2 μm. For the *syn*-4mer/LC8 specimen, a dataset of 34 micrographs was collected and picked in an automated fashion to select the center of ~4 – 5 nm densities, corresponding to individual LC8 dimers, using DoG Picker (55). We note that a variety of alternative automated particle picking tools were assayed for this workflow (35, 50, 51), which included traditional blob-pickers, template-based methods, as well as neural net particle picking algorithms. Following this initial screen, DoG Picker was selected due to the ease-of-use and performance as compared to these alternative methods.

DoG Picker settings were optimized for radius equal to 8 pixels and optimal thresholds ranging from 4.0 – 4.4, resulting in ~2000 – 3700 particle picks per micrograph with minimal contribution from background, assessed manually. From this dataset, 4 micrographs were set aside for training that contained a total of 14,306 LC8 particles. A separate validation set of 5 micrographs was prepared similarly using DoG Picker, yielding 17,245 particles. A dataset of 104 micrographs of the Nup159 construct were collected under identical conditions, that yielded a total of 246,328 particles by automated selection using DoG Picker. For the *d*ASCIZ construct, our previously collected dataset of 305 micrographs was re-processed and used for automated analysis (22), and yielding 557,134 LC8 particles by DoG Picker.

For use in method development and validation studies, the *syn-*4mer training dataset was curated by the microscopist familiar with the LC8-IDP structure to manually classify a representative set of LC8 oligomers as 2-mers, 3-mers, 4-mers, etc. To minimize ambiguity, the microscopist selected complexes that were well separated from neighboring particles on the micrograph. This procedure resulted in a curated set of 54 oligomers of varying valency (containing 216 LC8 particles in total) that were used for calibration of our automated analysis workflow.

For further comparative analysis, additional single-particle datasets were obtained by manual selection from the recorded micrographs for traditional 2D classification and averaging in EMAN (35). Obtained image stacks contained 4151 putative oligomers extracted with a box size of 96 for the LC8/*syn*-4mer dataset, 5875 oligomers and a box size of 96 for the LC8/Nup159 dataset, and 2434 complexes with a box size of 160 for LC8/*d*AZCIS dataset.

### Automated identification and population counting of oligomers

The automated pipeline for identifying beads-on-a-string LC8/IDP oligomers employed three stages: (i) clustering (ii) oligomer identification based on a scoring function (Fig. 2E,F), and (iii) distance-filtering to disregard crowded regions of micrographs (Supplemental Fig. 5). Single-linkage clustering (56) of all LC8 coordinates from the auto-picked micrographs was first performed. In this clustering method, data points separated by less than a given distance are grouped together to distinguishing sets of particles that cannot form an oligomer based on their inter-particle coordinates. The linkage distance was set to a value that is larger separation of neighboring LC8 binding sites on the IDP, as derived by the distribution of separation distances obtained in the curated training set (see Supplemental Figs. 3, 7, 9). In particular, the clustering threshold was set to 7 nm for the 4-site IDP, 8 nm for the Nup159 system, and 15 nm for the ASCIZ system.

A scoring algorithm was developed to classify the heterogeneous oligomer populations. The scoring algorithm is informed by the particle intensity (*I*), *i.e.*, the average pixel value within a picked particle as reported by the DoG picker (55) and the oligomer geometry, *i.e.*, particle-to-particle separation distance, (*d*) and angle (*Θ*) defined by three adjoining particles. The oligomer obtained from the hand curated training data were used to calibrate these features from their distributions in the training data. The distributions of these three metrics (Supplemental Fig. 4) provided parameters to score new oligomers, as described in equation (1).

To treat all three metrics *I, d*, and *Θ* on an equal basis, we used their training data cumulative distribution functions (CDFs) within their 0.5 to 99.5 percentile region. To give a higher score to small distances and angles between two or three given particles, we used 1-CDF as the probability scores *P_d_* and *P_θ_* for these two metrics, whereas the CDF was used as the probability score *P_I_* for the intensity *I* of a given particle. The total score for any *n*-mer is the normalized sum over all of its sequential intensity, distance, and angle log-probability scores:

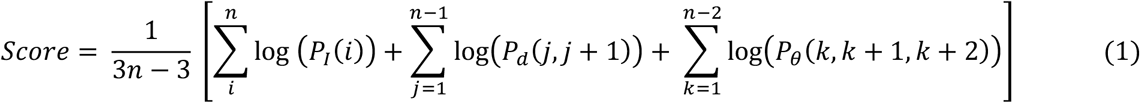

Here P = CDF or 1-CDF as noted above.

To obtain oligomer assignments from the clustered particles, our program considers every possible combination of particle sequences (or oligomeric states) within a cluster and scores them independently. The potential oligomers were then ranked by their length and their total score, thus giving precedence to longer assemblies (regardless of score) over shorter ones, and the highest-ranked non-overlapping oligomers in each cluster were saved. Finally, a score threshold value was applied to discard oligomers with a low total score, for instance such oligomers that consist of several low-intensity particles and with geometry that is unfavorable compared to the training set statistics. Based on Supplemental Fig. 4, a small (*i.e.*, more permissive) threshold value of 0.05 was selected for the analysis of LC8/*syn*-4mer and LC8/Nup159 and, based on Supplemental Fig. 10, a threshold value of 0.3 was applied to the analysis of LC8/*d*ASCIZ, which contains a significantly longer IDP.

In order to prevent assignments in crowded, ambiguous regions of the micrograph, oligomer assignments were filtered by a distancing criterion, counting only those oligomers that were separated at least by a specified distance from any other LC8 particles. For consistency with manual evaluation, an initial set of micrographs with automatically assigned oligomers at different distance thresholds were examined by a microscopist to identify an optimal value for this filtering distance. The threshold was set to 9 nm for the *syn-*4mer and Nup159 systems and to 14 nm for the *d*ASCIZ system. The corresponding fractional populations of all three systems are not significantly different at slightly larger or smaller filtering distance thresholds (+/- 2 nm) as shown in Supplemental Fig. 5. To facilitate direct comparison of oligomeric state populations from the corresponding manually and automatically assigned micrographs, the same threshold was applied to the manual dataset of *syn-*4mer and *d*ASCIZ systems.

### Correcting oligomer populations through self-consistent statistical re-scoring

Ultimately, the accuracy of oligomer prediction depends not only on the correct assignment of oligomers but also on the identification of artifactual structures that should not be counted – *i.e.*, spurious oligomers resulting from random proximity of free LC8 particles not bound to any IDP. Random proximity would extend actual *n*-mers to be wrongly counted as (*n+1*)-mers or longer. Recall that the IDPs “strings” themselves are not visible in the micrographs. To provide an estimate for the actual number of the underlying real oligomers, the experimental process of random placement of single LC8 particles was iteratively simulated and the degree of artifactual oligomer creation was evaluated in order to obtain a self-consistent estimate of the underlying populations.

The iterative correction procedure is initialized by randomly relocating all free LC8 particles, *i.e.*, those that were not assigned to be part of a putative oligomer during the initial identification and scoring process. This population of free particles are then positioned randomly and independently, but with a minimum distance of 2 nm from any other particle present on the micrograph, which roughly corresponds to the minimum distance of LC8 particles observed experimentally. This procedure produces a synthetic micrograph that includes all predicted oligomers from the original experimental micrograph, but with the single LC8s rearranged. Applying the same scoring and counting rules described above to this synthetic micrograph leads to different oligomer assignments due to the random relocation of free LC8 particles that can lead to the appearance of both new oligomer creations (randomly placed LC8s that meet our scoring criteria of an oligomer) and putative oligomer extensions (randomly placed LC8s that are now located near the terminus of a previously assigned *n*-mer). Another effect of this process is that oligomers that were previously counted in the experimental micrograph may now be “disqualified” because of the distancing criterion that is applied (to avoid assignments in crowded regions). At the same time, other oligomers which did not meet the distancing criteria in the original assignment process, because they were “blocked” by a nearby free LC8 particle(s), may now be “released” and counted.

The process just described is iterated until self-consistency is obtained between population counts from the synthetic micrographs, as compared to the originally assigned micrograph. Specifically, at each iteration *i*, the naive count of each *n-*mer oligomer species (abbreviated by *n*) in the synthetic micrograph is compared with that in the experimental micrograph to obtain the difference Δ_*i*_(*n*). If at iteration *k*, the cumulative sum of these differences over all previous iterations was a positive integer, i.e., 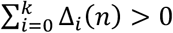, suggesting that the number of directly counted *n*-mers in the given synthetic micrograph exceeded those in the experimental micrograph, then that many putative *n*-mers were selected at random and pruned. Pruning of oligomers was performed by stripping one of their terminal particles and adding it to the set of free particles at iteration (*k*+1), thereby also reducing the number of putative *n*-mers and increasing that of putative (*n-1*)-mers. This operation was performed at every iteration in a cascading fashion from longer oligomers to shorter oligomers and the 2-mers were pruned by splitting and adding both particles to the set of free LC8s. If 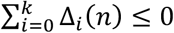, then no pruning and updating of putative oligomer counts was performed. This iterative process was conducted until the counts of all oligomer species in the synthetic micrograph matched those in the experimental micrograph. At that point, the updated population of putative oligomers can be considered corrected with respect to artifacts arising from the large number of free LC8 particles. If continued, the populations fluctuate among a set of values consistent with the originally assigned dataset.

Convergence was reached within 100 iterations for the *syn-*4mer system and within 200 iterations for the Nup159 and *d*ASCIZ systems (Fig. 3 and Supplemental Figs. 7,9). For each system, the naïve and corrected oligomer populations were obtained as the arithmetic mean over the last 50 iterations and the total population counts over all analyzed micrographs were converted into fractional populations. Error bars are derived as the square-root of the sum of all per-micrograph variances computed from the last 50 iterations, representing the effective standard deviation – *i.e.*, the scale of variation of the obtained mean values from the correction procedure. A flowchart describing the statistical correction procedure is provided in Supplemental Fig. 6.

## Code and data availability

All codes are available at https://github.com/ZuckermanLab/EM_OligomerAnalysis. Electron microscopy images and particle coordinate files are available at http://doi.org/10.5281/zenodo.4726027. Expression vectors are available upon request to E.B.

## Acknowledgments and Funding Sources

We thank Patrick Reardon for his assistance with collecting the AUC data, and the staff at the OHSU Multiscale Microscopy Core for assistance and training. The research was funded by the National Science Foundation (MCB 1715823 to D.M.Z.) and (MCB 1617019 to E.B.) and the National Institutes of Health (R35GM124779 and R01EY030987 to S.L.R.).

## Author Contributions

All authors contributed to the preparation of the manuscript. B.M. and R.M contributed equally. B.M. developed the single-particle distribution and statistical analysis programs. R.M. conducted the electron microscopy experiments and analysis. A.E. and J.H. performed the biochemistry and biophysical characterizations of LC8/IDP complexes. E.B. supervised the biochemical and biophysical studies. S.L.R. supervised the electron microscopy studies. D.M.Z. supervised the theoretical modeling and statistical analysis. E.B., S.L.R. and D.M.Z. provided overall oversight to the design and execution of the work.

## Competing Interest Statement

The authors have no competing interests.

## SUPPLEMENTAL INFORMATION

### (SUPPLEMENTAL FIGURES, TABLES AND LEGENDS)

#### SUPPLEMENTAL FIGURES AND LEGENDS

**Supplemental Fig. 1:**
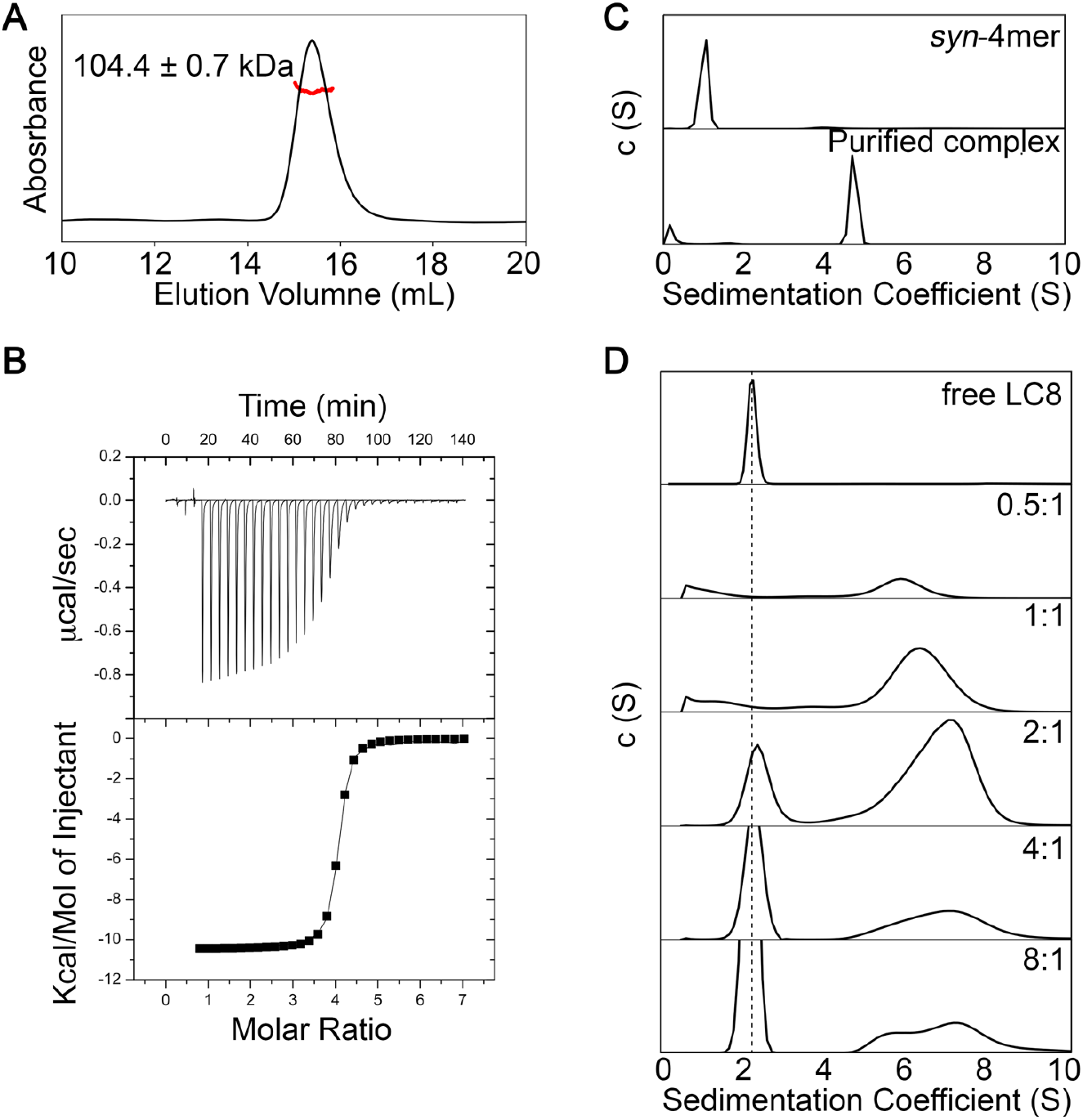
Sedimentation velocity analytical ultracentrifugation (AUC) of LC8 complexes. A) SEC-MALS of *syn*-4mer in complex with LC8. Purified complex eluted as a single peak, with a mass of 104.4±0.7 kDa, within uncertainty of the expected mass for a 2:8 complex, 105.2 kDa. B) Isotherm of binding between LC8 and the syn-4mer. The isotherm fits well to a simple binding model with K_d_ = 36±3 nM, ΔH = 10.47±0.04 kcal/mol, and N=3.98±0.01. Model fit is shown as a line. C) AUC data for the syn-4mer and size exclusion purified LC8/syn-4mer complex. A sharp peak at a sedimentation coefficient of 4.7 S indicates a tight and homogeneous complex. D) AUC data for LC8 and LC8/Nup159 complexes formed at increasing ratios of LC8. The dashed line is centered on the LC8 peak. The multiple peaks in the 6-8 S for the complex indicates heterogeneity of the complex and with an S value close to 8, it suggests a higher order assembly than a 5-mer and two Nup159 chains.

**Supplemental Fig. 2:**
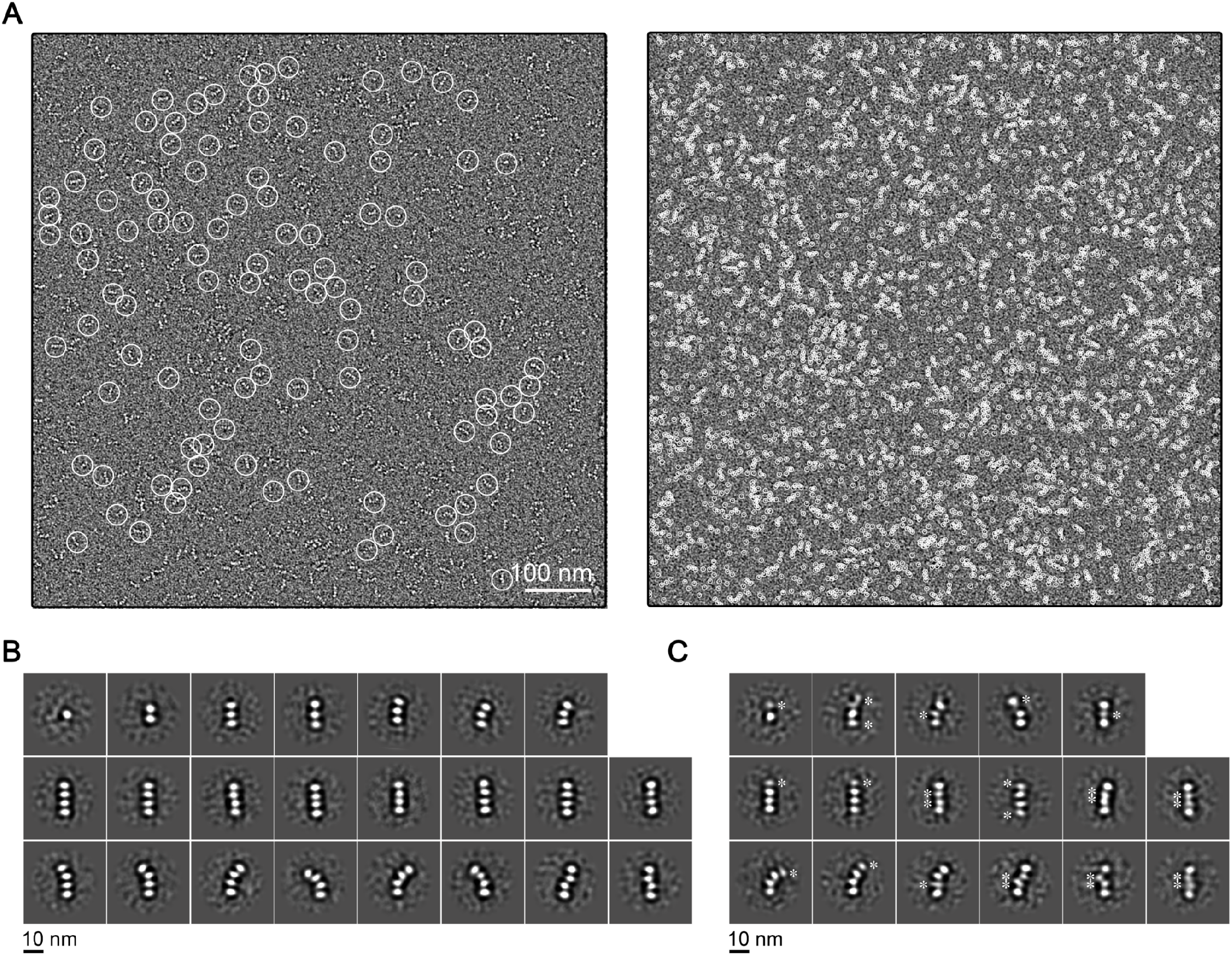
Single-particle EM analysis of LC8/*syn*-4mer. A) Representative micrograph of LC8/*syn*-4mer showing (*left*) manually selected oligomers and (*right*) auto-picked LC8 densities using DoG Picker (55). Selected particles are indicated with white circles. Scale bar = 100 nm. B) Expanded set of two-dimensional (2D) class-averages depicting free LC8 (*top left*) and a range of assembled *syn*-4mer/LC8 complexes (2-mers to 4-mers) present in the image dataset that were well-resolved. C) Expanded set of 2D class-averages showing assembled *syn*-4mer/LC8 complexes displaying varying degree of conformational heterogeneity. Asterisk indicate densities of LC8 that display blurred features that are less well-resolved, indicative of unresolved conformational/configurational heterogeneity. Scale bar = 10 nm in panels B and C.

**Supplemental Fig. 3:**
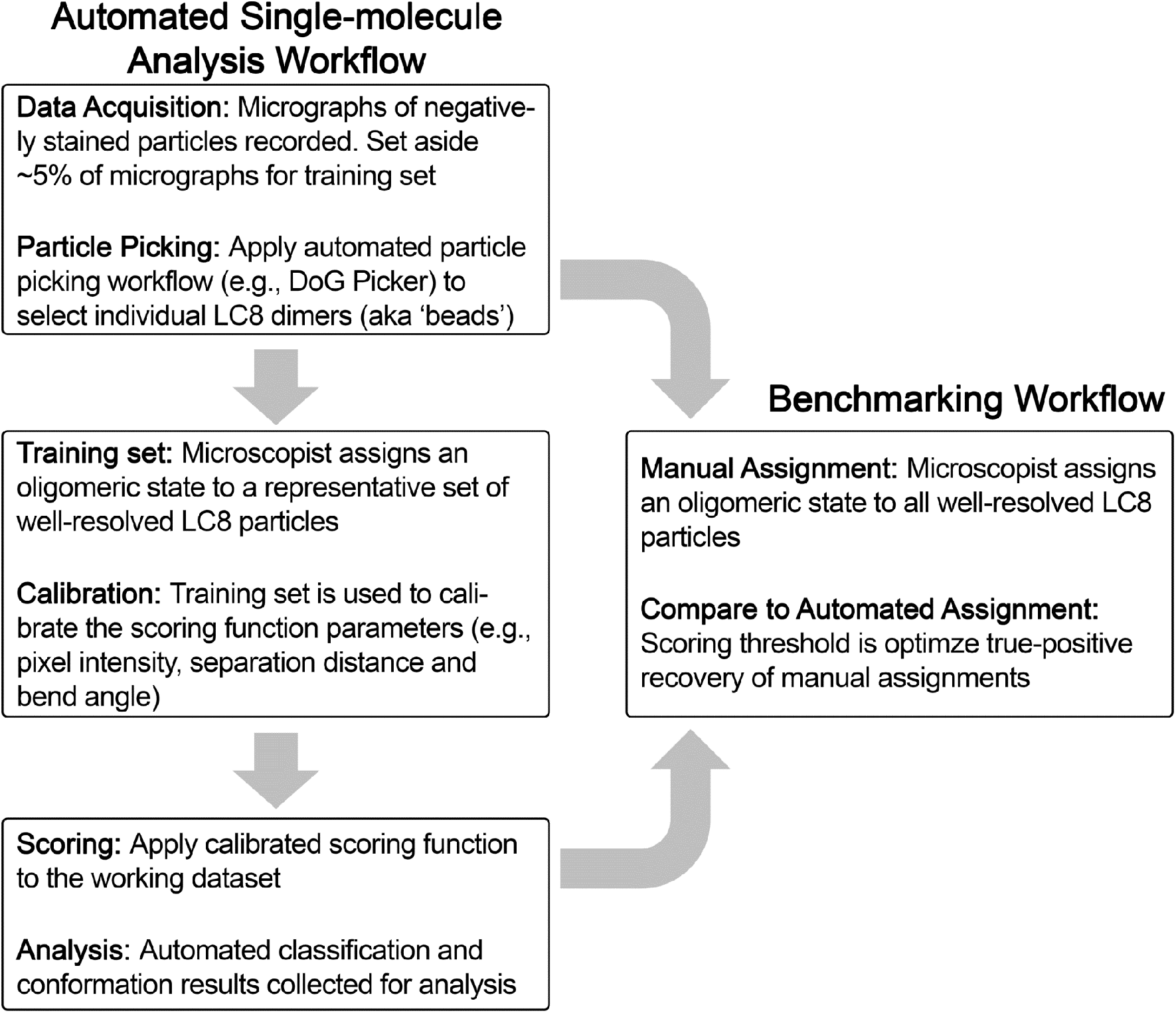
Automated single-particle distribution analysis and benchmarking workflow. Diagram depicting the individual steps of the automated oligomer assignment and benchmarking procedures. The Analysis Workflow was applied to the LC8/*syn-*4mer, the LC8/Nup159, and the LC8/*d*ASCIZ datasets. The Benchmarking Workflow was applied to the LC8/*syn-*4mer system to validate the results.

**Supplemental Fig. 4:**
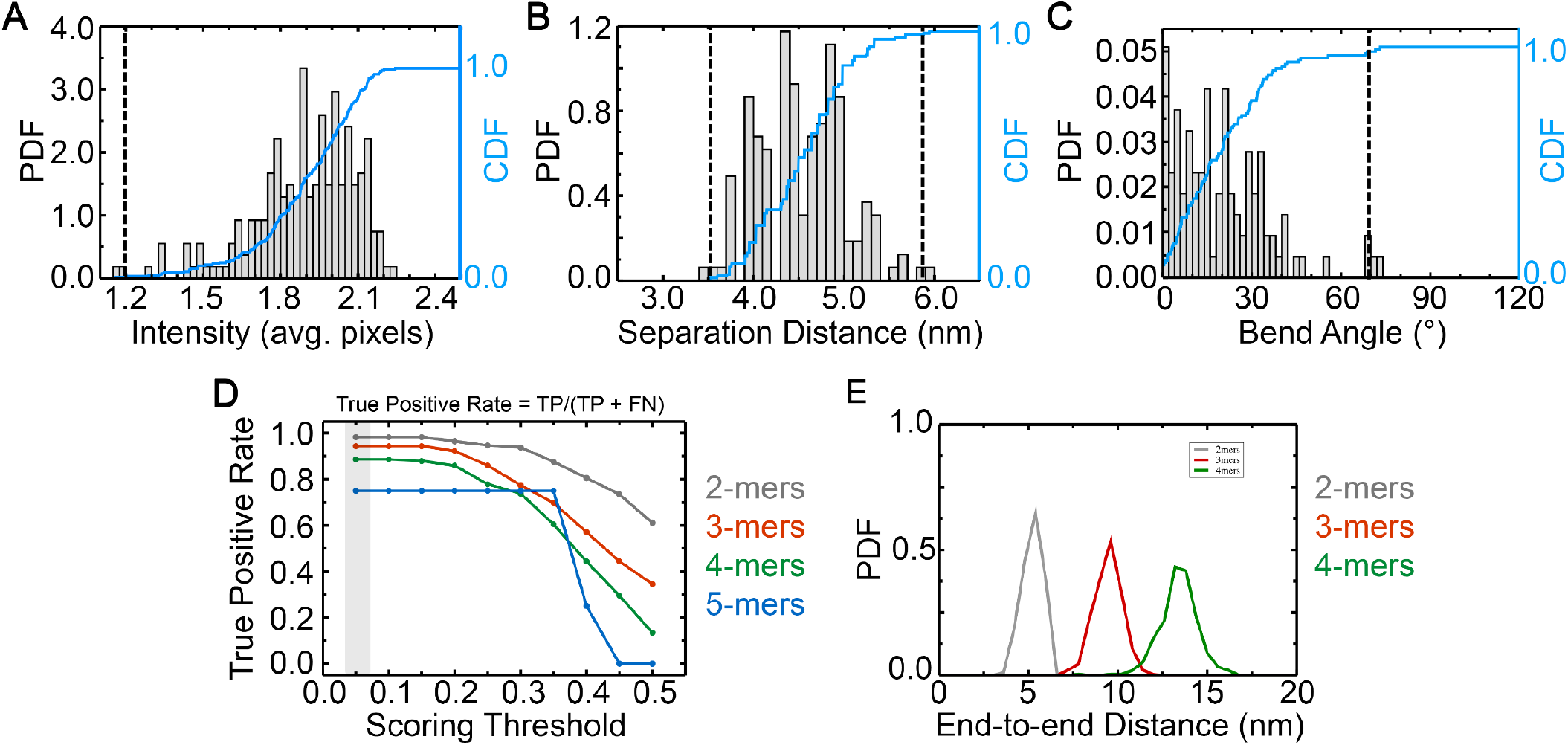
Scoring function calibration for the syn-4mer/LC8 complex. A – C) Distributions of (A) LC8 particle intensities (n = 216), (B) LC8-to-LC8 distance separations (*d*) (n = 162) and (C) bend angles (n = 108) between sequential LC8-LC8 ‘bond’ vectors (θ), obtained from a training dataset of syn-4mer/LC8 micrographs. The probability density function (PDF, gray bars) is shown together with the cumulative distribution function (CDF, blue). Outlier values with CDF < 0.005 and/or > 0.995 were excluded for calibration of the scoring function and given a score of zero, as indicated by the vertical dashed lines. D) The True Positive Rate, defined as the fraction of manually assigned oligomers that were also automatically assigned, for 2-mers (grey), 3-mers (red), 4-mers (green), and 5-mers (blue). A smaller (more permissive) threshold value = 0.05 yielded the most accurate automatic assignments, and was selected for application to the full dataset (transparent grey). E) Distribution of end-to-end distances, *i.e.*, the distance between terminal LC8s in an oligomer, based on all oligomers predicted for this system. Data displayed independently for 2-mers (gray, n= 1312), 3-mers (red, n= 485) and 4-mer (green, n= 359).

**Supplemental Fig. 5:**
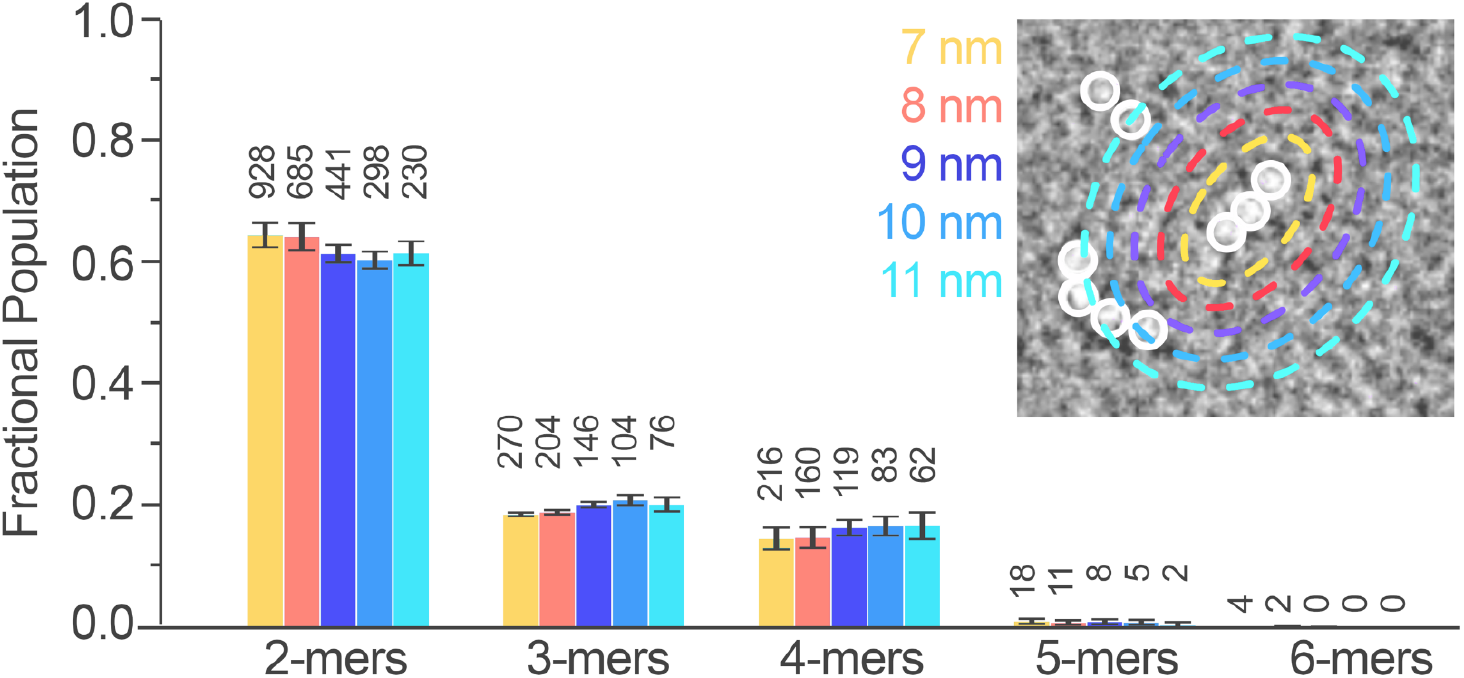
Effects of distance thresholding on assigned oligomer populations. The fractional population distribution of oligomeric states automatically assigned with different values for the nearest neighbor distance criterion, applied to a training set of LC8/*syn*-4mer particles. The numbers on top of bars indicate total population counts. *Inset*, zoom of representative region of an electron micrograph depicting an assigned oligomer in a crowded region of neighboring particles (white circles), with distance radii indicated as dotted circles. The closest distance from any oligomer in the reference oligomer (inset, center) to any other LC8 particle is considered.

**Supplemental Fig. 6:**
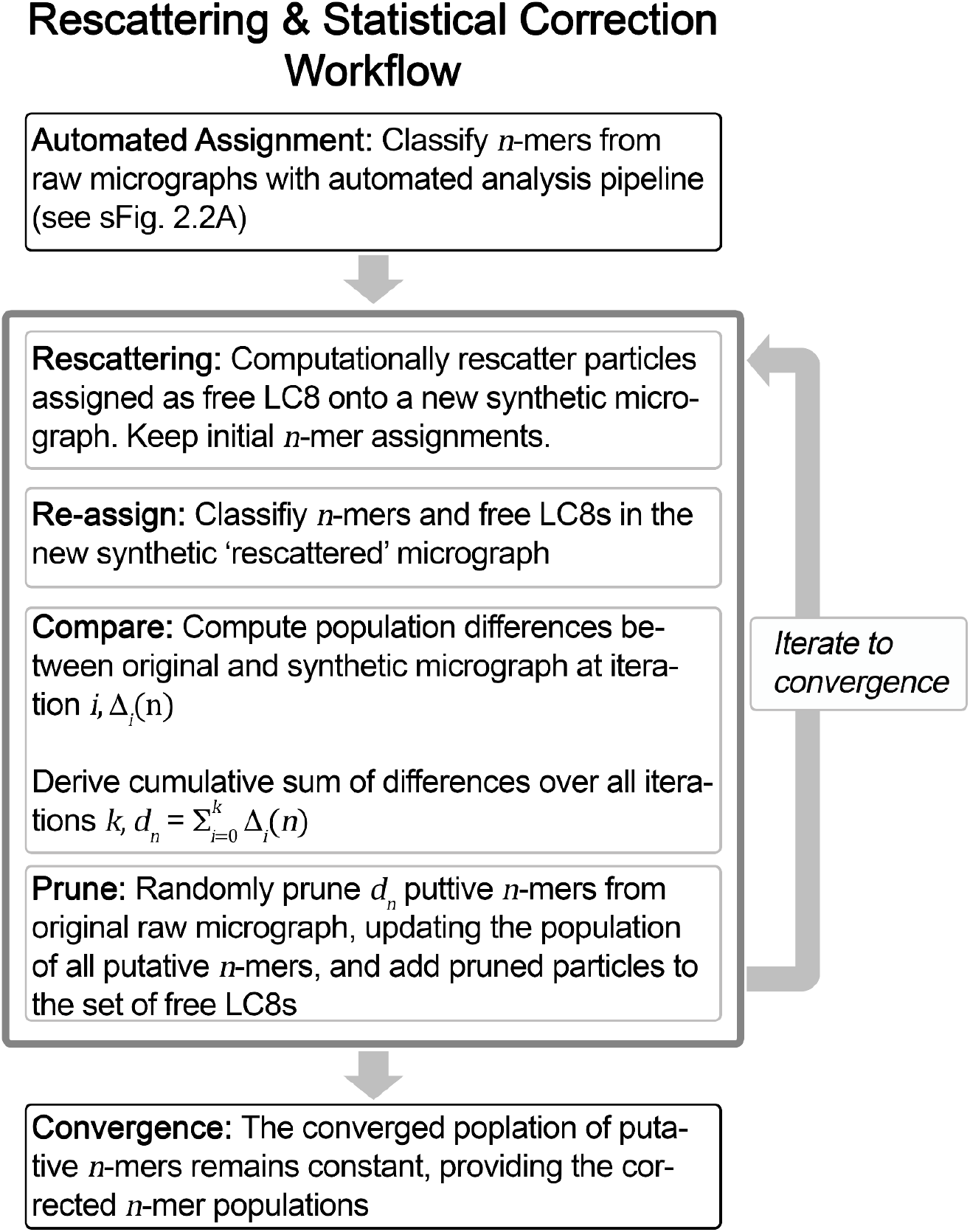
Flow-chart of the re-scattering and statistical re-scoring protocol. Diagram depicting the individual steps of the statistical correction procedure summarized in Fig. 3A.

**Supplemental Fig. 7:**
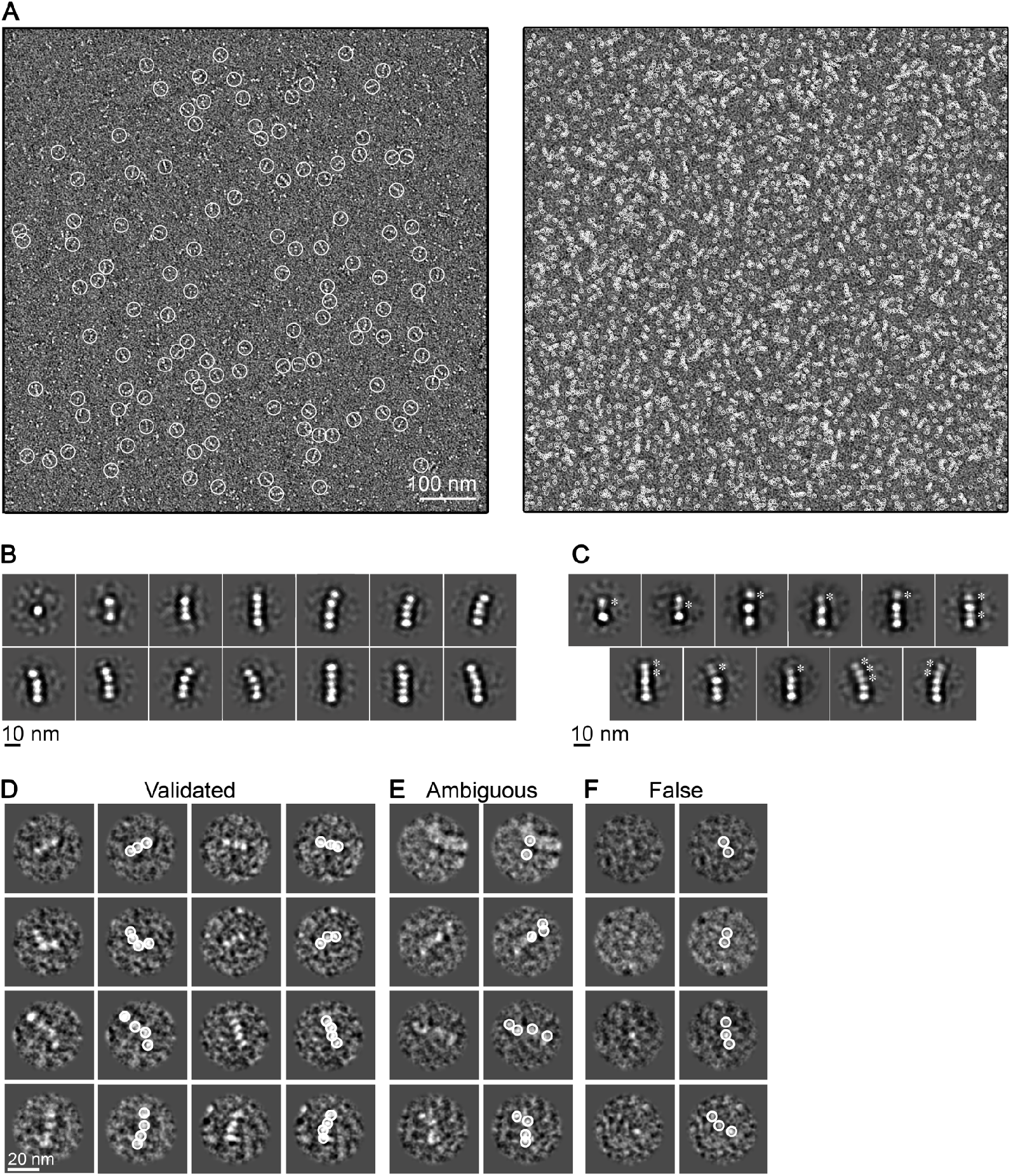
Single-particle EM analysis of LC8/Nup159. A) Representative micrograph of LC8/Nup159 showing (*left*) manually selected oligomers and (*right*) auto-picked LC8 densities using DoG Picker (55). Selected particles are indicated with white circles. Scale bar = 100 nm. B) Expanded set of representative two-dimensional (2D) class-averages depicting free LC8 (top left) and a range of assembled LC8/Nup159 complexes (2-mers to 5-mers) present in the image dataset that were well-resolved. C) Expanded set of 2D class-averages showing assembled LC8/Nup159 complexes displaying varying degree of conformational heterogeneity. Asterisk indicate densities of LC8 that display blurred features that are less well-resolved, indicative of unresolved conformational/configurational heterogeneity. Scale bar = 10 nm in panels B and C. D - F) Microscopist validation of automated single-particle assignments. D) Representative images of auto-assigned LC8/Nup159 oligomers that were deemed acceptable by the microscopists. E) Representative images of complexes assigned by scoring function and deemed to be too ambiguous to confidently assign by the microscopists upon evaluation (*e.g.*, containing weak LC8 density and/or neighboring LC8 densities that were not autopicked). F) Representative images of complexes assigned by the scoring function and deemed to be too falsely assigned by the microscopist upon re-evaluation (*e.g.*, containing one or more autopicked densities corresponding to background carbon). For panels D – F, raw images shown in the left column and autopicked results obtained by DoG Picker shown by white circles. Scale bar = 20 nm.

**Supplemental Fig. 8:**
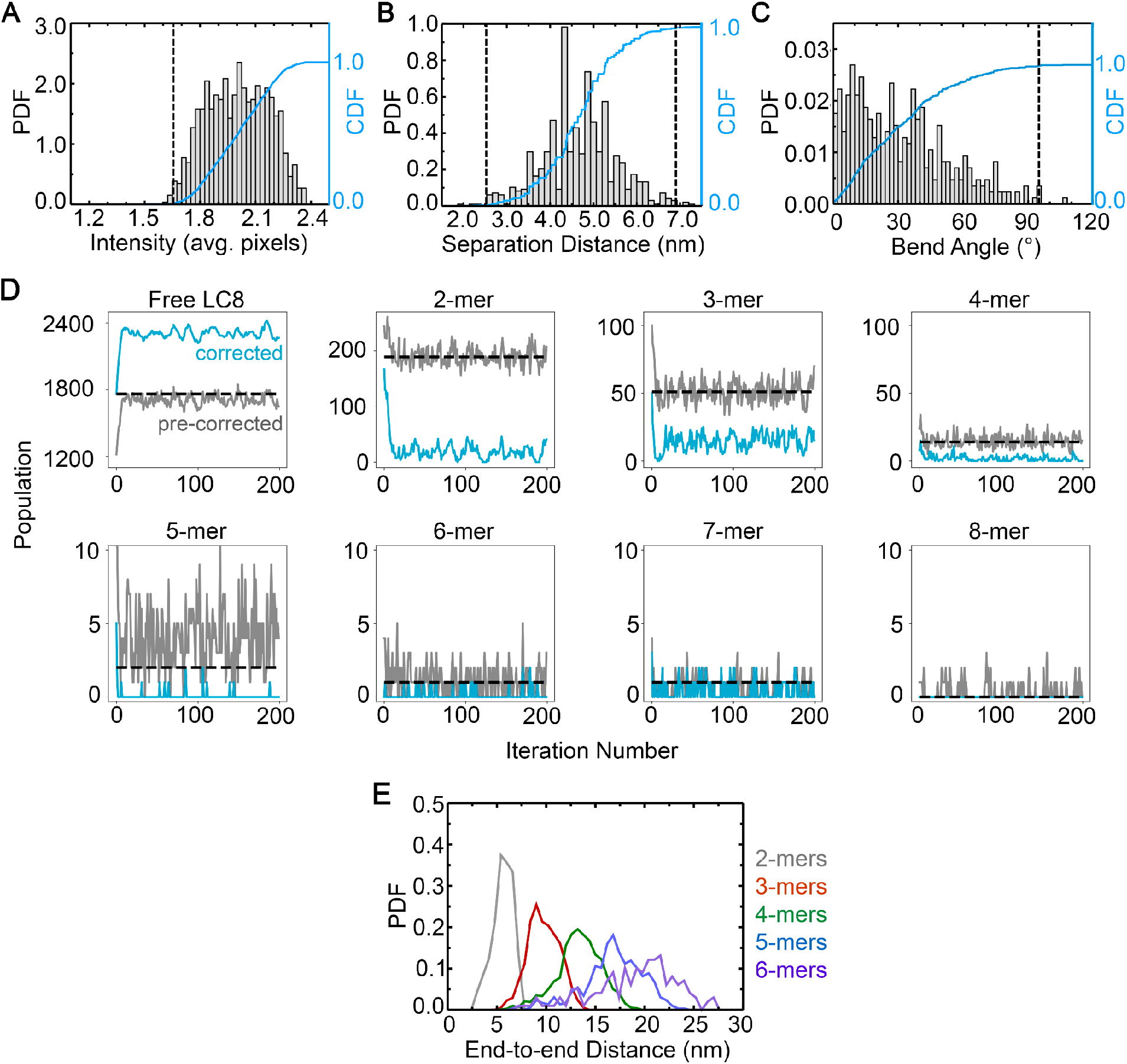
Scoring function calibration, statistical correction and end-to-end distance analysis of Nup159/LC8. A – C) Distributions of (A) LC8 particle intensities (n = 1037), (B) LC8-to-LC8 distance separations (n = 732) (*d*) and (C) bend angles (n = 427) between sequential LC8-LC8 ‘bond’ vectors (θ), obtained from a training dataset of Nup159/LC8 micrographs. The probability density function (PDF, gray bars) is shown together with the cumulative distribution function (CDF, blue). Outlier values with CDF < 0.005 and/or > 0.995 were excluded for calibration of the scoring function and given a score of zero, as indicated by the vertical dashed lines. D) The pre-corrected (gray) and corrected (cyan) populations of oligomers and free LC8 as a function of iteration number during the statistical correction simulation of one example micrograph. The dashed black lines indicate the corresponding populations of initial oligomer assignments in the experimental micrograph, to which the gray lines converge. E) Distribution of end-to-end distances, *i.e.*, the distance between terminal LC8s in an oligomer, based on all oligomers predicted for this system. Data displayed independently for 2-mers (gray, n= 9786), 3-mers (red, n= 3083), 4-mer (green, n= 1196), 5-mers (blue, n= 489), and 6-mers (purple, n= 165).

**Supplemental Fig. 9:**
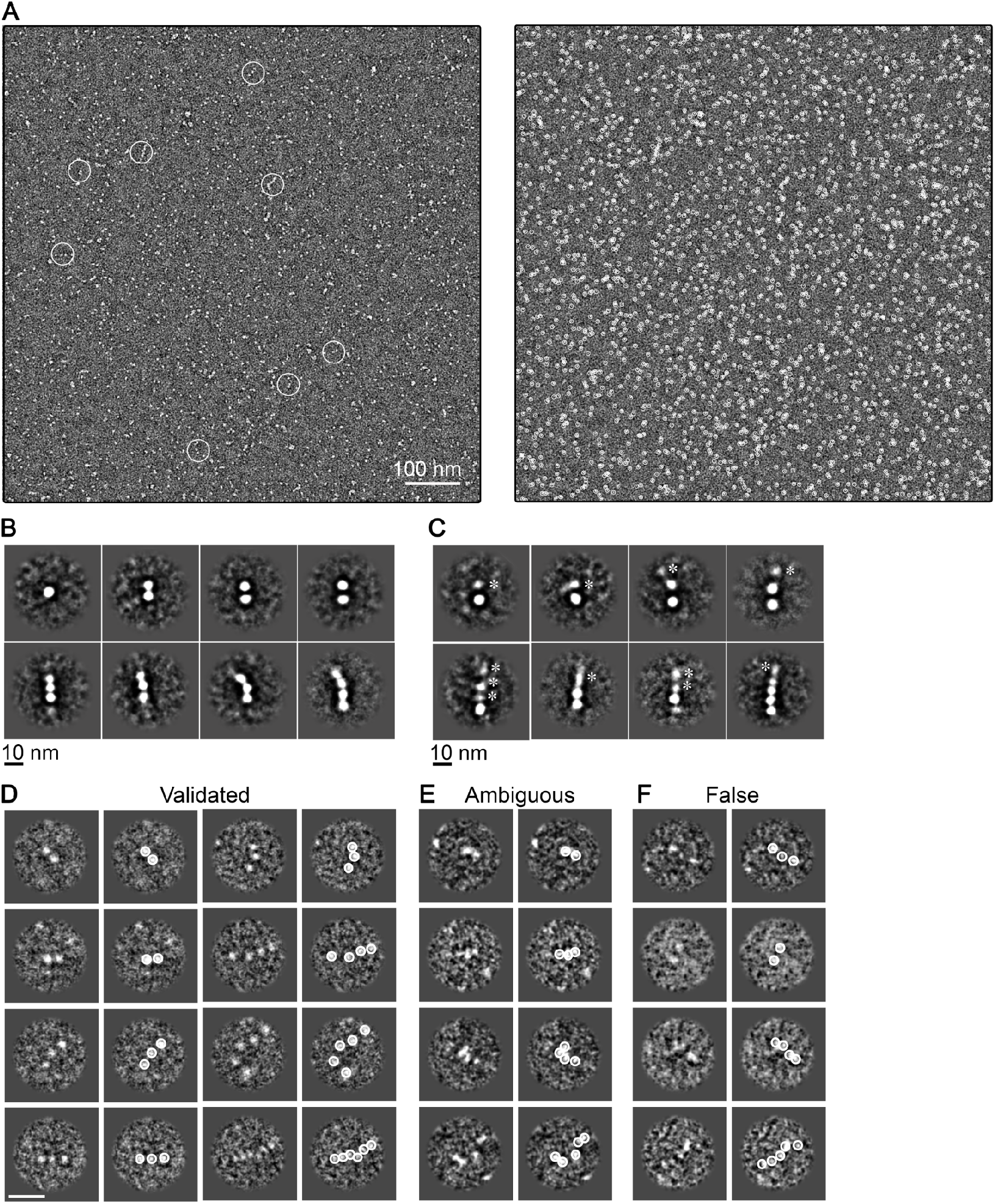
Single-particle EM analysis of LC8/*d*ASCIZ. A) Representative micrograph of LC8/*d*ASCIZ showing (*left*) manually selected oligomers and (*right*) auto-picked LC8 densities using DoG Picker (55). Selected particles are indicated with white circles. Scale bar = 100 nm. B) Expanded set of 2D class-averages depicting free LC8 (top left) and a range of assembled *d*ASCIZ/LC8 complexes (2-mers to 4-mers) present in the image dataset that were well-resolved. C) Expanded set of 2D class-averages showing assembled *d*ASCIZ/LC8 complexes displaying varying degree of conformational heterogeneity. Asterisk indicate densities of LC8 that display blurred features that are less well-resolved, indicative of unresolved conformational/configurational heterogeneity. Scale bar = 10 nm in panels B and C. D – F) Microscopist validation of automated single-particle assignments. D) Representative images of auto-assigned LC8/*d*ASCIZ oligomers that were deemed acceptable by the microscopists. E) Representative images of complexes assigned by scoring function and deemed to be too ambiguous to confidently assign by the microscopists upon evaluation (*e.g.*, containing weak LC8 density and/or neighboring LC8 densities that were not autopicked). F) Representative images of complexes assigned by the scoring function and deemed to be too falsely assigned by the microscopist upon re-evaluation (*e.g.*, containing one or more autopicked densities corresponding to background carbon). For panels D – F, raw images shown in the left column and autopicked results obtained by DoG Picker shown by white circles. Scale bar = 20 nm.

**Supplemental Fig. 10:**
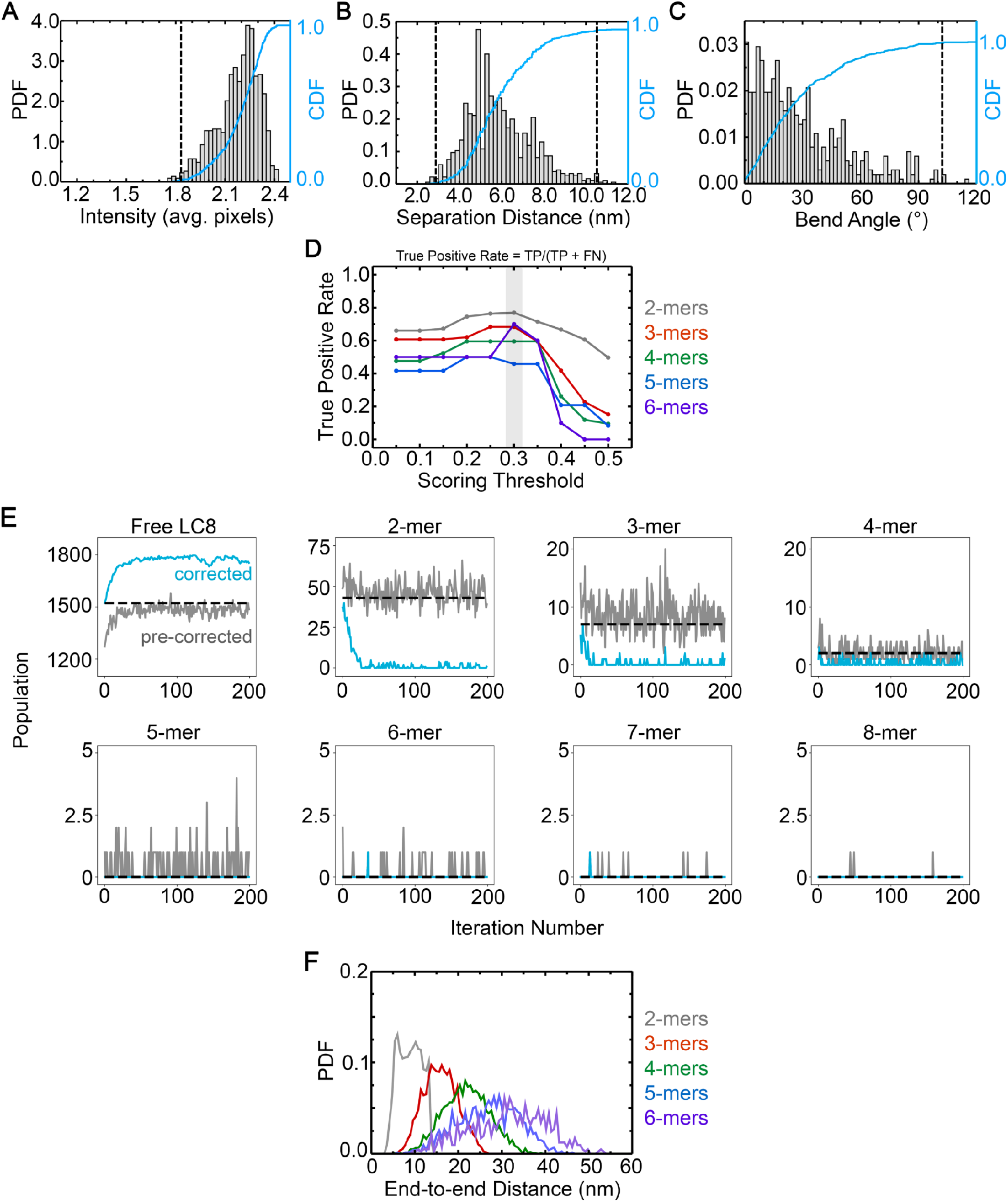
Scoring function calibration, statistical correction and end-to-end distance analysis of *d*ASCIZ/LC8. A – C) Distributions of (A) LC8 particle intensities (n = 1360), (B) LC8-to-LC8 distance separations (*d*) (n = 925) and (C) bend angles between sequential LC8-LC8 ‘bond’ vectors (θ) (n = 509), obtained from a training dataset of syn-4mer/LC8 micrographs. The probability density function (PDF, gray bars) is shown together with the cumulative distribution function (CDF, blue). Outlier values with CDF < 0.005 and/or > 0.995 were excluded for calibration of the scoring function and given a score of zero, as indicated by the vertical dashed lines. D) The True Positive Rate, i.e., the fraction of manually assigned oligomers that were also automatically assigned, for 2-mers (gray), 3-mers (red), 4-mers (green), 5-mers (blue), and 6-mers (purple). A threshold value of 0.3 yields the most accurate automatic assignments and was selected for application to the full dataset (transparent grey). E) The pre-corrected (gray) and corrected (cyan) populations of oligomers and free LC8 as a function of iteration number during the statistical correction simulation of one example micrograph. The dashed black lines indicate the corresponding populations of initial oligomer assignments in the experimental micrograph, to which the gray lines converge. F) Distribution of end-to-end distances, *i.e.*, the distance between terminal LC8s in an oligomer, based on all oligomers predicted for this system. Data displayed independently for 2-mers (grey, n = 27823), 3-mers (red, n = 6136), 4-mer (green, n = 1591), 5-mers (blue, n = 530), and 6-mers (purple, n = 215).

### SUPPLEMENTAL TABLES

**Supplemental Table 1:** Overview of EM data collection and parameters.

|  | *syn*-4mer | Nup159 | *dASCIZ* |
| --- | --- | --- | --- |
| # micrographs | 30 | 104 | 305 |
| Defocus range (μm) | 1.9–4.0 | 1.6–4.0 | 2.0–3.5 |
| Pixel size (Å/pixel) | 4.37 | 4.37 | 4.37 |
| Magnification | 49,000 | 49,000 | 49,000 |
| Microscope power (kV) | 120 | 120 | 120 |
| Picked complexes | 4151 | 5875 | 2434 |
| Picked LC8 domains | 14,333 | 246,328 | 557,134 |
| Dog-pick thresholds | 4; 1000 | 4; 1000 | 4; 1000 |
| Dog-pick radius (pixels) | 8 | 8 | 8 |

**Supplemental Table 2:**
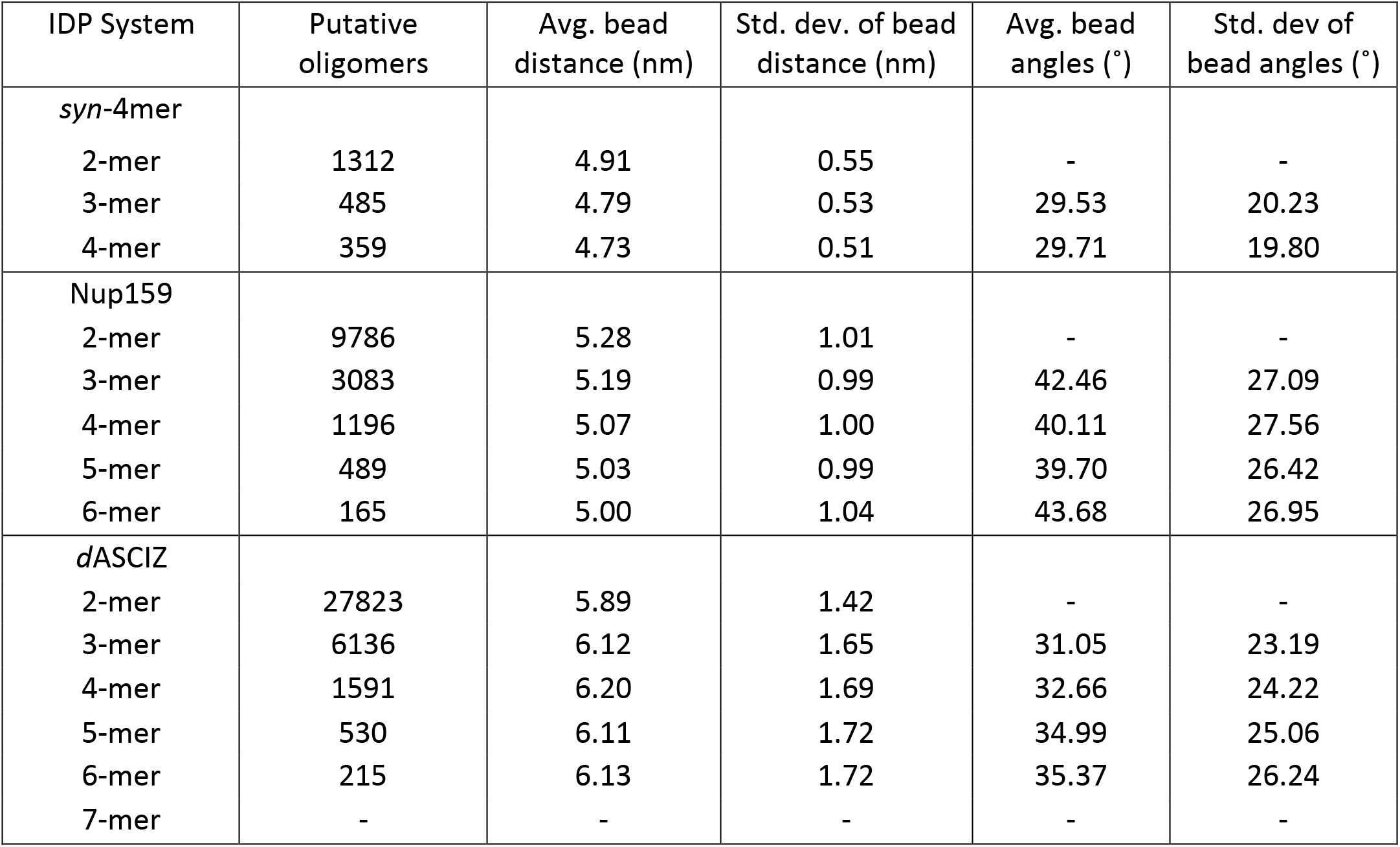
Overview of statistics obtained by singe particle distribution analysis.

| IDP System | Putative oligomers | Avg. bead distance (nm) | Std. dev. of bead distance (nm) | Avg. bead angles (°) | Std. dev of bead angles (°) |
| --- | --- | --- | --- | --- | --- |
| *syn*-4mer |  |  |  |  |  |
| 2-mer | 1312 | 4.91 | 0.55 | - | - |
| 3-mer | 485 | 4.79 | 0.53 | 29.53 | 20.23 |
| 4-mer | 359 | 4.73 | 0.51 | 29.71 | 19.80 |
| Nup159 |  |  |  |  |  |
| 2-mer | 9786 | 5.28 | 1.01 | - | - |
| 3-mer | 3083 | 5.19 | 0.99 | 42.46 | 27.09 |
| 4-mer | 1196 | 5.07 | 1.00 | 40.11 | 27.56 |
| 5-mer | 489 | 5.03 | 0.99 | 39.70 | 26.42 |
| 6-mer | 165 | 5.00 | 1.04 | 43.68 | 26.95 |
| *dASCIZ* |  |  |  |  |  |
| 2-mer | 27823 | 5.89 | 1.42 | - | - |
| 3-mer | 6136 | 6.12 | 1.65 | 31.05 | 23.19 |
| 4-mer | 1591 | 6.20 | 1.69 | 32.66 | 24.22 |
| 5-mer | 530 | 6.11 | 1.72 | 34.99 | 25.06 |
| 6-mer | 215 | 6.13 | 1.72 | 35.37 | 26.24 |
| 7-mer | - | - | - | - | - |

### SUPPLEMENTAL MOVIE LEGEND

**Supplemental Movie 1**

**Single-particle conformational heterogeneity in the *syn*-4mer/LC8 complex.** Montage of selected syn-4mer/LC8 particles aligned and oriented to display the conformational heterogeneity observed captured from the raw image dataset by the automated assignment procedure. Individual LC8 dimers are highlighted by a white.

**Supplemental Movie 2**

**Annotated conformational heterogeneity in the *syn*-4mer/LC8 complex.** Annotated version of syn-4mer/LC8 particles aligned and oriented as in Figure 2F. Individual LC8 dimers are indicated by a grey circles.

## Notes

### Competing Interest Statement

The authors have declared no competing interest.

